# Ancient exapted transposable elements promote nuclear enrichment of human long noncoding RNAs

**DOI:** 10.1101/189753

**Authors:** Joana Carlevaro-Fita, Taisia Polidori, Monalisa Das, Carmen Navarro, Tatjana I. Zoller, Rory Johnson

**Affiliations:** Department for BioMedical Research (DBMR), University of Bern, 3008 Bern, Switzerland; Department of Medical Oncology, Inselspital, University Hospital and University of Bern, 3010 Bern, Switzerland; Graduate School of Cellular and Biomedical Sciences, University of Bern, Bern, Switzerland; Department of Computer Science and Artificial Intelligence, University of Granada, Spain

**Keywords:** Transposable element, subcellular localization, long noncoding RNA, lncRNA, evolution, exaptation

## Abstract

The sequence domains underlying long noncoding RNA (lncRNA) activities, including their characteristic nuclear enrichment, remain largely unknown. It has been proposed that these domains can originate from neofunctionalised fragments of transposable elements (TEs), otherwise known as RIDLs (Repeat Insertion Domains of Long Noncoding RNA), although just a handful have been identified. It is challenging to distinguish functional RIDL instances against a numerous genomic background of neutrally-evolving TEs. We here show evidence that a subset of TE types experience evolutionary selection in the context of lncRNA exons. Together these comprise an enrichment group of 5374 TE fragments in 3566 loci. Their host lncRNAs tend to be functionally validated and associated with disease. This RIDL group was used to explore the relationship between TEs and lncRNA subcellular localisation. Using global localisation data from ten human cell lines, we uncover a dose-dependent relationship between nuclear/cytoplasmic distribution, and evolutionarily-conserved L2b, MIRb and MIRc elements. This is observed in multiple cell types, and is unaffected by confounders of transcript length or expression. Experimental validation with engineered transgenes shows that these TEs drive nuclear enrichment in a natural sequence context. Together these data reveal a role for TEs in regulating the subcellular localisation of lncRNAs.

## Introduction

The human genome contains many thousands of long noncoding RNAs (lncRNAs), of which at least a fraction are likely to have evolutionarily-selected biological functions (Ulitsky and Bartel 2013). Our current working hypothesis is that, similar to proteins, lncRNA functions are encoded in primary sequence through “domains”, or discrete elements that mediate specific aspects of lncRNA activity. Such activities range from molecular interactions to subcellular localisation (Guttman and Rinn 2012; Mercer and Mattick 2013; Johnson and Guigó 2014). Experimental support for this domain model is beginning to emerge (Marín-Béjar et al. 2017). Mapping domains in a comprehensive manner is thus a key step towards the understanding and prediction of lncRNA functions.

One possible source of lncRNA domains are transposable elements (TEs) (Johnson and Guigó 2014). TEs are known to have been major contributors to genomic evolution through the insertion and neofunctionalisation of sequence fragments – a process known as *exaptation* (Bourque 2009)(Feschotte 2008). This process has contributed to the evolution of diverse features in genomic DNA, including transcriptional regulatory motifs (Bourque et al. 2008; Johnson et al. 2006), microRNAs (Roberts et al. 2014), gene promoters (Faulkner et al.; Huda et al. 2011), and splice sites (Lev-Maor et al. 2003; Sela et al. 2007).

We recently proposed that exaptation also takes place in the context of lncRNAs, with TEs contributing pre-formed functional domains. We termed these “RIDLs” – *Repeat Insertion Domains of Long noncoding RNAs* (Johnson and Guigó 2014). As RNA, TEs are known to interact with a rich variety of proteins, meaning that in the context of lncRNA they could plausibly act as protein-docking sites (Blackwell et al. 2012). Diverse evidence also points to repetitive sequences forming intermolecular Watson-Crick RNA:RNA and RNA:DNA hybrids (Johnson and Guigó 2014; Gong and Maquat 2011; Holdt et al. 2013). However, it is likely that *bona fide* RIDLs represent a small minority of the many exonic TEs, with the remainder being phenotypically-neutral “passengers”.

A small but growing number of RIDLs have been described, reviewed in (Johnson and Guigó 2014). These are found in lncRNAs with clearly-demonstrated functions, including the X-chromosome silencing transcript *XIST* (Elisaphenko et al. 2008), the oncogene *ANRIL* (Holdt et al. 2013) and the regulatory antisense *UchlAS* (Carrieri et al. 2012). In each case, domains of repetitive origin are necessary for a defined function: the structured A-repeat of *XIST*, of retroviral origin, recruits the PRC2 silencing complex (Elisaphenko et al. 2008); Watson-Crick hybridisation between RNA and DNA *Alu* elements recruits *ANRIL* to target genes (Holdt et al. 2013); a SINEB2 repeat in *Uchl1AS* increases translational rate of its sense mRNA (Carrieri et al. 2012). In parallel, transcriptome-wide maps of lncRNA-linked TEs have shown how TEs have contributed extensively to lncRNA gene evolution (Kelley and Rinn 2012; Kapusta et al. 2013)(Schmitt et al. 2016)(Hezroni et al. 2015). However, there has been no attempt to enrich these maps for RIDLs with evidence of selected functions in the context of mature lncRNA molecules.

Subcellular localisation, and the domains controlling it, are crucial determinants of lncRNA functions (reviewed in (Chen 2016)). For example, transcriptional regulatory lncRNAs must be located in the nucleus and chromatin, whereas those regulating microRNAs or translation should be present in the cytoplasm (Zhang et al. 2014b). Although higher nuclear/cytoplasmic ratios are a hallmark of lncRNAs, a large population of cytoplasmic transcripts also exists (Mukherjee et al. 2017) (Carlevaro-Fita et al. 2016; Derrien et al. 2012)(Cabili et al. 2015; Mas-Ponte et al. 2017)(Benoit Bouvrette et al. 2018). If lessons learned from mRNA are also valid for lncRNAs, then short sequence motifs recognised by RNA binding proteins (RBPs) will be an important localisation-regulatory mechanism (Martin and Ephrussi 2009). This was recently demonstrated for the *BORG* lncRNA, where a pentameric motif was shown to mediate nuclear retention (Zhang et al. 2014a). Similarly, multiple copies of the 156 bp RRD repeat motif mediate nuclear enrichment of the *FIRRE* lncRNA, through binding to hnRNPU (Hacisuleyman et al. 2016a) (Hacisuleyman et al. 2014). Another study implicated an inverted pair of *Alu* elements in nuclear retention of *lincRNA-P21* (Chillón and Pyle 2016). This raises the possibility that, by “copying and pasting” generic RNA motifs, RIDLs could fine-tune lncRNA localisation at a global scale.

The aim of the present study is to create a human transcriptome-wide catalogue of putative RIDLs. Supporting its relevance, lncRNAs carrying these RIDLs are enriched for functional genes. Finally, we provide *in silico* and experimental evidence that certain RIDL types, derived from ancient transposable elements, promote the nuclear enrichment of their host transcripts.

## Results

The objective of this study is to create a map of repeat insertion domains of long noncoding RNAs (RIDLs) and link them to lncRNA functions. We hypothesise that RIDLs could confer such functions through interactions with DNA, RNA or protein molecules (Johnson and Guigó 2014) (Figure 1A).

**Figure 1:**
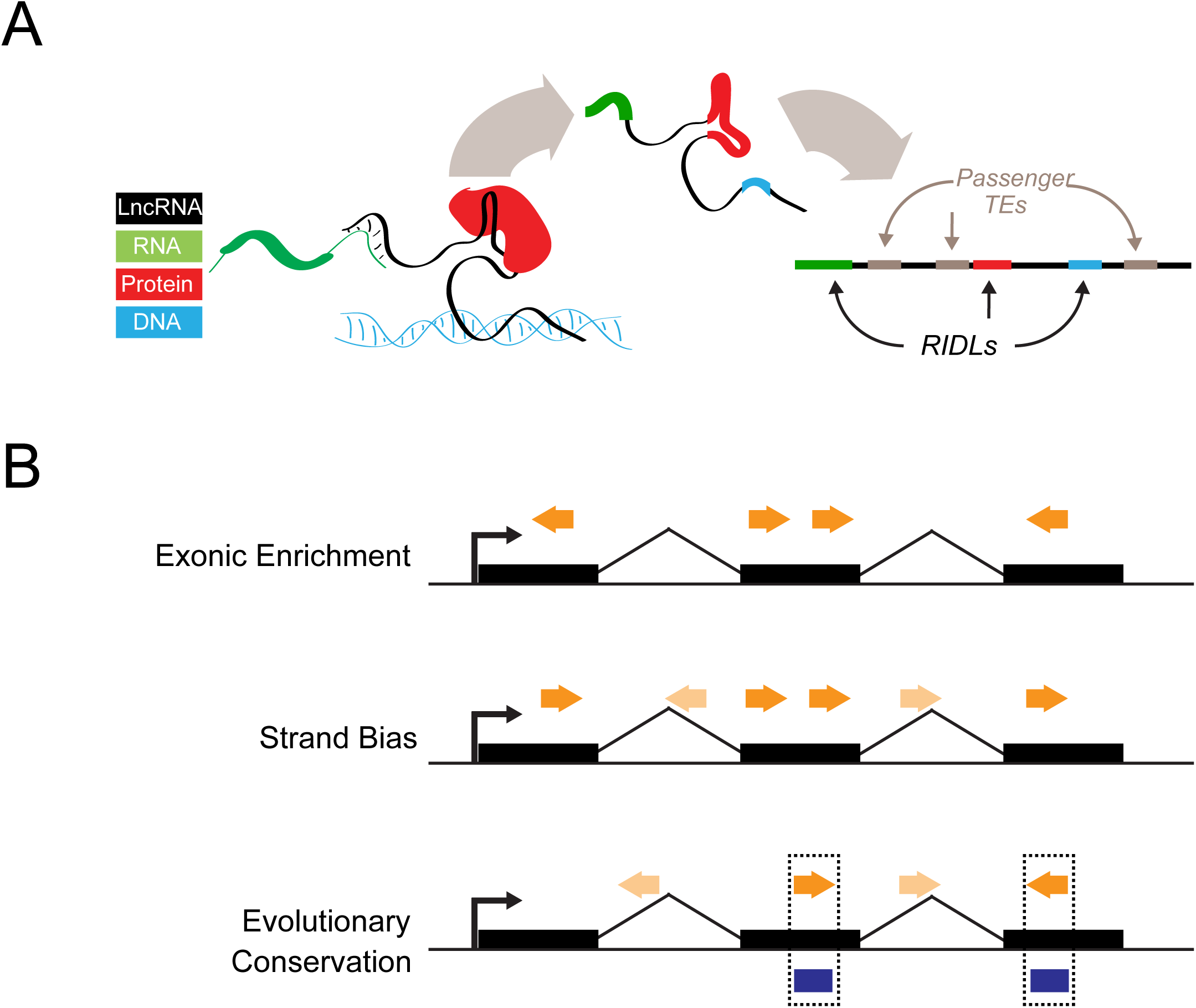
Repeat insertion domains of lncRNAs (RIDLs). (A) In the “Repeat Insertion Domain of LncRNAs” (RIDL) model, exonically-inserted fragments of transposable elements contain pre-formed protein-binding (red), RNA-binding (green) or DNA-binding (blue) activities, that contribute to functionality of the host lncRNA (black). RIDLs are likely to be a small minority of exonic TEs, coexisting with large numbers of non-functional “passengers” (grey). (B) RIDLs (dark orange arrows) will be distinguished from passenger TEs by signals of selection, including: (1) simple enrichment in exons; (2) a preference for residing on a particular strand relative to the host transcript; (3) elevated evolutionary conservation in exons compared introns. Selection might be identified by comparing exonic TEs to a neutral population, for example those residing in lncRNA introns (light coloured arrows).

Any attempt to map RIDLs must deal with two challenges. First, that they will likely represent a small minority amongst many phenotypically-neutral “passenger” transposable elements (TEs) in lncRNA exons (Figure 1B). Second, many TE instances may be under evolutionary selection, but for functions executed at the *DNA level* (eg transcription factor binding sites, enhancer elements), rather than the RNA level (Bassett et al. 2014)

Therefore, it is necessary to identify RIDLs by some signature of selection that is specific for a mature RNA product using an appropriate background model. In this study we use three types of such signatures: exonic enrichment, strand bias (with respect to host gene), and exon-specific evolutionary conservation (Figure 1B). To estimate background, we utilise intronic TEs, since they should mirror any biases of TE distribution across the genome but are not incorporated into mature lncRNA transcripts.

Resulting RIDL predictions should be considered as “enrichment groups”, due to high rates of false positive predictions, and all downstream analyses should be interpreted accordingly.

### A map of exonic transposable elements in GENCODE v21 lncRNAs

Our first aim was to create a comprehensive map of transposable elements (TEs) within the exons of GENCODE v21 human lncRNAs (Figure 2A). Altogether 5,520,018 distinct TE insertions were intersected with 48684 exons from 26414 transcripts of 15877 GENCODE v21 lncRNA genes, resulting in 46474 exonic TE insertions in lncRNA (Figure 1B). 13121 lncRNA genes (82.6%) carry at least one exonic TE fragment in one or more of their mature transcripts.

**Figure 2:**
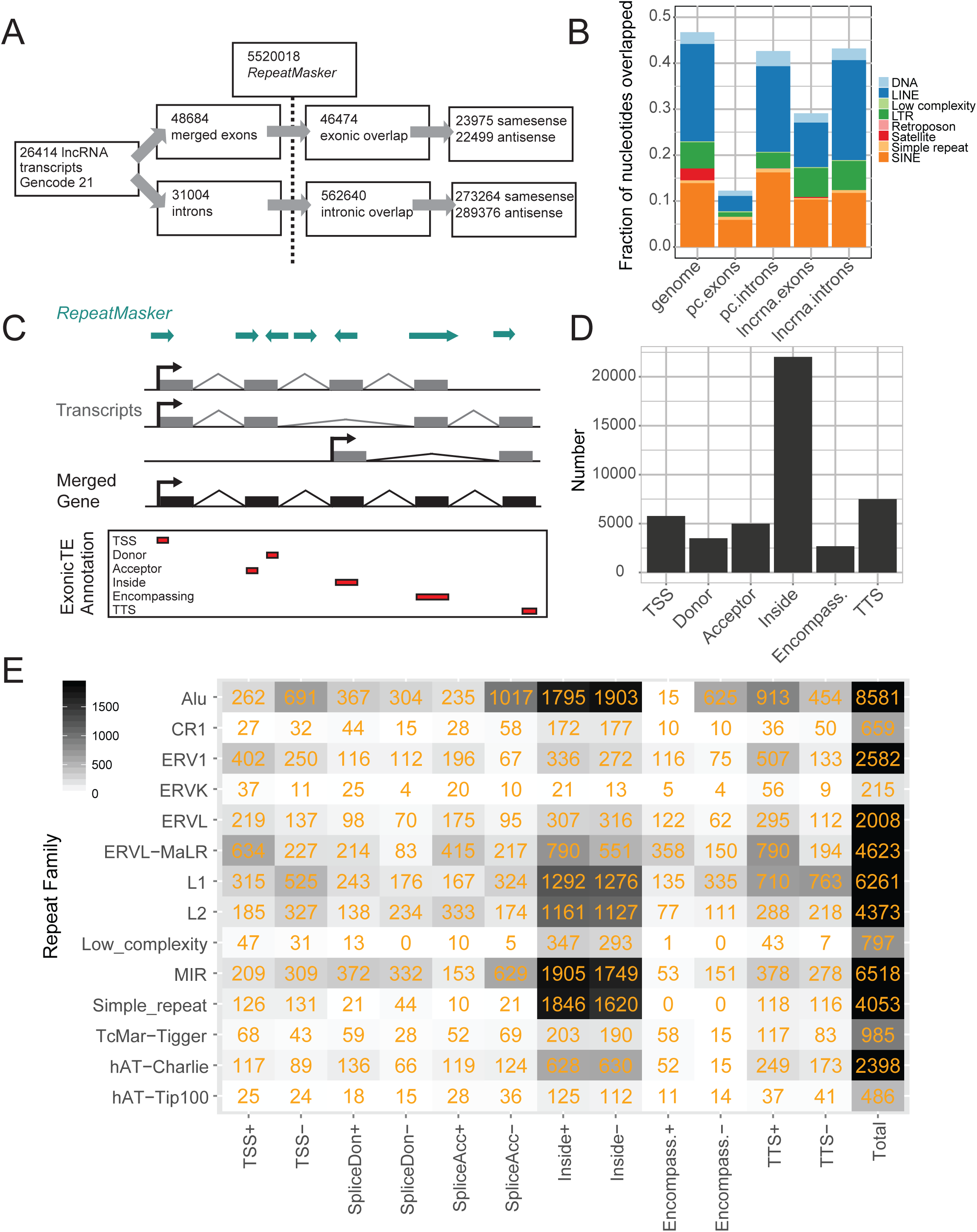
An exonic transposable element annotation with the GENCODE v21 lncRNA catalogue. (A) Statistics for the exonic TE annotation process using GENCODE v21 lncRNAs. (B) The fraction of nucleotides overlapped by TEs for lncRNA exons and introns, protein-coding introns and exons (“pc”), and the whole genome. (C) Overview of the annotation process. The exons of all transcripts within a lncRNA gene annotation are merged. Merged exons are intersected with the RepeatMasker TE annotation. Intersecting TEs are classified into one of six categories (bottom panel) according to the gene structure with which they intersect, and the relative strand of the TE with respect to the gene: “TSS”, overlapping the transcription start site; “Donor”, splice donor site; “Acceptor”, splice acceptor site; “Inside”, the TE boundaries both lie within the exon; “Encompassing”, the exon boundaries both lie within the TE; “TTS”, the transcription termination site. (D) Summary of classification breakdown for exonic TE annotation. (E) Classification of TE classes in exonic TE annotation. Numbers indicate instances of each type. +/-indicate the relative strand of the TE with respect to lncRNA transcript.

We also created a reference dataset with 31,004 GENCODE lncRNA introns, resulting in 562,640 intron-overlapping TE fragments (Figure 2A). Comparing intronic and exonic TE data, we see that lncRNA exons are depleted for TE insertions: 29.2% of exonic nucleotides are of TE origin, compared to 43.4% of intronic nucleotides (Figure 2B), similar to previous studies (Kapusta et al. 2013). This may reflect generalised selection against disruption of functional lncRNA transcripts by TEs. The exonic depletion of TEs in lncRNAs is less pronounced than for protein-coding loci, whereas the intronic TE density of both is similar to the whole-genome average.

### Contribution of transposable elements to lncRNA gene structures

TEs have contributed widely to both coding and noncoding gene structures by the insertion of elements such as promoters, splice sites and termination sites (Sela et al. 2007). We next classified inserted TEs by their contribution to lncRNA gene structure (Figure 2C,D). It should be borne in mind that this analysis is dependent on the accuracy of underlying GENCODE annotations, which are often incomplete at 5′ and 3′ ends (Lagarde et al. 2017). Altogether 4993 (18.9%) transcripts’ promoters lie within a TE, most often those of *Alu*, L1 and ERVL-MaLR classes (Figure 2E). 7497 (28.4%) lncRNA transcripts are terminated by a TE, most commonly by L1, *Alu*, ERVL-MaLR classes. 8494 lncRNA splice sites (32.2%) are of TE origin, and 2681 entire exons are fully contributed by TEs (10.1%) (Figure 2E). These observations support known contributions of TEs to gene structural features (Sela et al. 2007). Nevertheless, the most frequent case is represented by 22,031 TEs that lie completely within an exon and do not overlap any splice junction (“inside”).

### Evidence for selection on certain exonic transposable element types

This exonic TE map represents the starting point for the identification of RIDLs, defined as the subset of TEs with evidence for functionality in the context of mature lncRNAs. In this and subsequent analyses, TEs are grouped by type as defined by *RepeatMasker*. We utilise three distinct sources of evidence for selection on TEs: exonic enrichment, strand bias and evolutionary conservation (Figure 1B).

We first asked whether particular TE types are enriched in lncRNA exons, compared to intronic sequence (Kelley and Rinn 2012). Thus, we calculated the ratio of exonic / intronic sequence coverage by TEs (Figure 3A). We found enrichment >2-fold for numerous repeat types, including endogenous retrovirus classes (HERVE-int, HERVK9-int, HERV3-int, LTR12D) in addition to others such as ALR/Alpha, BSR/Beta and REP522. A number of simple repeats are also enriched in lncRNA, including GC-rich repeats. A weaker but more generalized trend of enrichment is also observed for various MLT repeat classes. These findings are consistent with previous analyses by Kelley and Rinn using whole genome, rather than introns, as background (Kelley and Rinn 2012). Similarly, both studies agree in finding no difference in *Alu* density between lncRNA exons and intergenic / intronic DNA.

**Figure 3:**
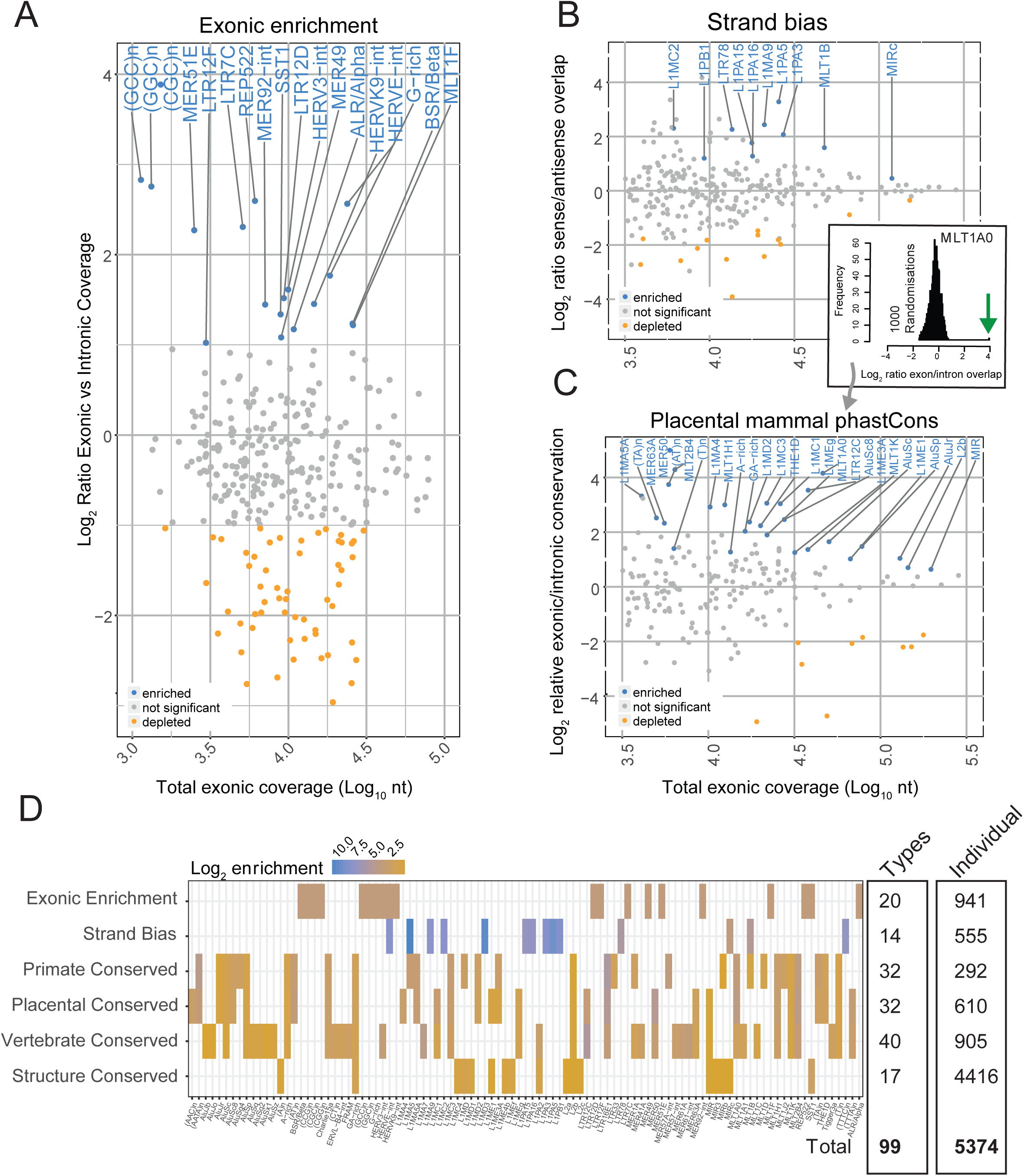
Evidence for selection on transposable elements in lncRNA exons. (A) Figure shows, for every TE type, the enrichment of per nucleotide coverage in exons compared to introns (*y* axis) and overall exonic nucleotide coverage (*x* axis). Enriched TE types (at a 2-fold cutoff) are shown in blue. (B) As for (A), but this time the *y* axis records the ratio of nucleotide coverage in sense vs antisense configuration. “Sense” here is defined as sense of TE annotation relative to the overlapping exon. Similar results for lncRNA introns may be found in Supplementary Figure S1. Significantly-enriched TE types are shown in blue. Statistical significance was estimated by a randomisation procedure, and significance is defined at an uncorrected empirical *p*-value < 0.001 (See Material and Methods).(C) As for (A), but here the *y* axis records the ratio of per-nucleotide overlap by phastCons mammalian-conserved elements for exons vs introns. Similar results for three other measures of evolutionary conservation may be found in Supplementary Figure S1. Significantly-enriched TE types are shown in blue. Statistical significance was estimated by a randomisation procedure, and significance is defined at an uncorrected empirical *p*-value < 0.001 (See Material and Methods). An example of significance estimation is shown in the inset: the distribution shows the exonic/intronic conservation ratio for 1000 simulations. The green arrow shows the true value, in this case for MLT1A0 type. (D) Summary of TE types with evidence of exonic selection. Six distinct evidence types are shown in rows, and TE types in columns. On the right are summary statistics for (i) the number of unique TE types identified by each method, and (ii) the number of instances of exonic TEs from each type with appropriate selection evidence. The latter are henceforth defined as “RIDLs”.

Despite their overall abundance throughout the genome, presently-active LINE1 elements are relatively depleted in lncRNA exons (Figure 3A). It is possible that this reflects selection against disruption to normal gene expression, where numerous weak polyadenylation signals lead to premature transcription termination when the LINE1 element lies on the same strand as the overlapping gene (Perepelitsa-Belancio and Deininger 2003). Other explanations may be low transcriptional processivity exhibited by the LINE1 ORF2 in the sense strand (Perepelitsa-Belancio and Deininger 2003), or else epigenetic silencing effects (Hollister and Gaut 2009).

As a second source of evidence for selection, we searched for TE types displaying a strand preference relative to host lncRNA (Johnson and Guigó 2014). We were conscious of a major source of bias: as shown above, many TSS and splice sites of lncRNA are contributed by TEs, and such cases would lead to artefactual strand bias. To avoid this, we ignored any TEs that overlap an exon-intron boundary. We calculated the relative strand overlap of all remaining TEs in lncRNA exons. Statistical significance was assessed by randomisation, with significance defined at P<0.001, corresponding to a false discovery rate (FDR) below 5% (similar cutoffs apply to subsequent analyses, more details may be found in Materials and Methods) (Figure 3B). In lncRNA exons, a number of TE types are enriched in either sense or antisense, dominated by LINE1 family members, possibly for the reasons mentioned above. Other significantly enriched TE types include LTR78, MLT1B, and MIRc (Figure 3B).

To test the specificity of this exonic strand bias, we performed equivalent analysis using introns. Although intronic strand bias is weaker, we did detect a modest yet statistically-significant depletion of same-strand TE insertions (Supplemental Figure S1). This is especially true for LINE1 elements, possibly for aforementioned reasons. In contrast to exons, almost no TE types were significantly enriched on the same-strand in introns.

To test for TE type-specific conservation, we turned to two sets of predictions of evolutionarily-conserved elements. First, the widely-used phastCons conserved elements, based on phylogenetic hidden Markov model (Siepel et al. 2005) calculated separately on primate, placental mammal and vertebrate alignments; second, the more recent “Evolutionarily Conserved Structures” (ECS) set (Smith et al. 2013). Importantly, the phastCons regions are defined based on sequence conservation alone, while the ECS are defined by phylogenetic analysis of RNA structure evolution.

To look for evidence of evolutionary conservation on exonic TEs, we calculated the fraction of nucleotides overlapped by evolutionarily-conserved genomic elements, and compared to the equivalent fraction for intronic TEs of the same type. To assess statistical significance, we again used positional randomisation (see inset in Figure 3C). This pipeline was applied independently to the phastCons (placental mammal shown in Figure 3C, primate and vertebrate in Supplemental Figure S1B,C) and ECS (Supplemental Figure S1D) data. The majority of TE types do not exhibit signatures of conservation (grey points). However, for each conservation type, the method detects significant conservation for a minority of TE types (Figure 3C). This enrichment disappeared when phastCons elements were positionally randomised (Supplemental Figure S2A). It is unlikely that overlap with protein-coding loci biases the results, since equivalent analyses using intergenic lncRNAs yielded similar candidate RIDLs (Supplemental Figure S2B). A similar analysis was performed using protein-coding exons, and although a number of significantly-conserved TEs were identified, they display limited overlap with those from lncRNAs (Supplemental Figure S2C). We also found a small number of TEs depleted for signatures of conservation in lncRNA exons, namely the young *Alu*Sz, *Alu*Sx and *Alu*Jb (phastCons) and L1M4c and *Alu*Sx1 (ECS) (coloured orange in Figure 3C and Supplemental Figure S1). The cause of this depletion is unclear, although one explanation is enrichment of conservation in intronic TEs due to RNA-independent regulatory roles as observed previously (Su et al. 2014).

All the selection evidence is summarised in Figure 3D. As might be expected, one observes a high degree of concordance in candidate TEs identified by the three phastCons methods, in addition to a smaller number with both phastCons and ECS evidence, including L2b and MIRb. This is not surprising given the distinct methodologies used to infer conservation. Less concordance is observed between conservation, enrichment, and strand bias candidates, although some TEs are identified by multiple methods, such as MIRc (strand bias and ECS).

### An annotation of RIDLs

We next combined all TE classes with evidence of functionality into a draft annotation of RIDLs. This annotation combined altogether 99 TE types with at least one type of selection evidence. For each TE / evidence pair, only those TE instances satisfying that evidence were included. In other words, if MIRb elements were found to be associated with vertebrate phastCons elements, then *only* those instances of exonic MIRb elements overlapping such an element would be included in the RIDL annotation, and all other exonic MIRs would be excluded. This operation was performed for all three phastCons element types, ECS elements and strand-bias. An example is *CCAT1* lncRNA oncogene: it carries three exonic MIR elements, of which one is defined as a RIDL based on its overlapping a phastCons element (Figure 4A).

**Figure 4:**
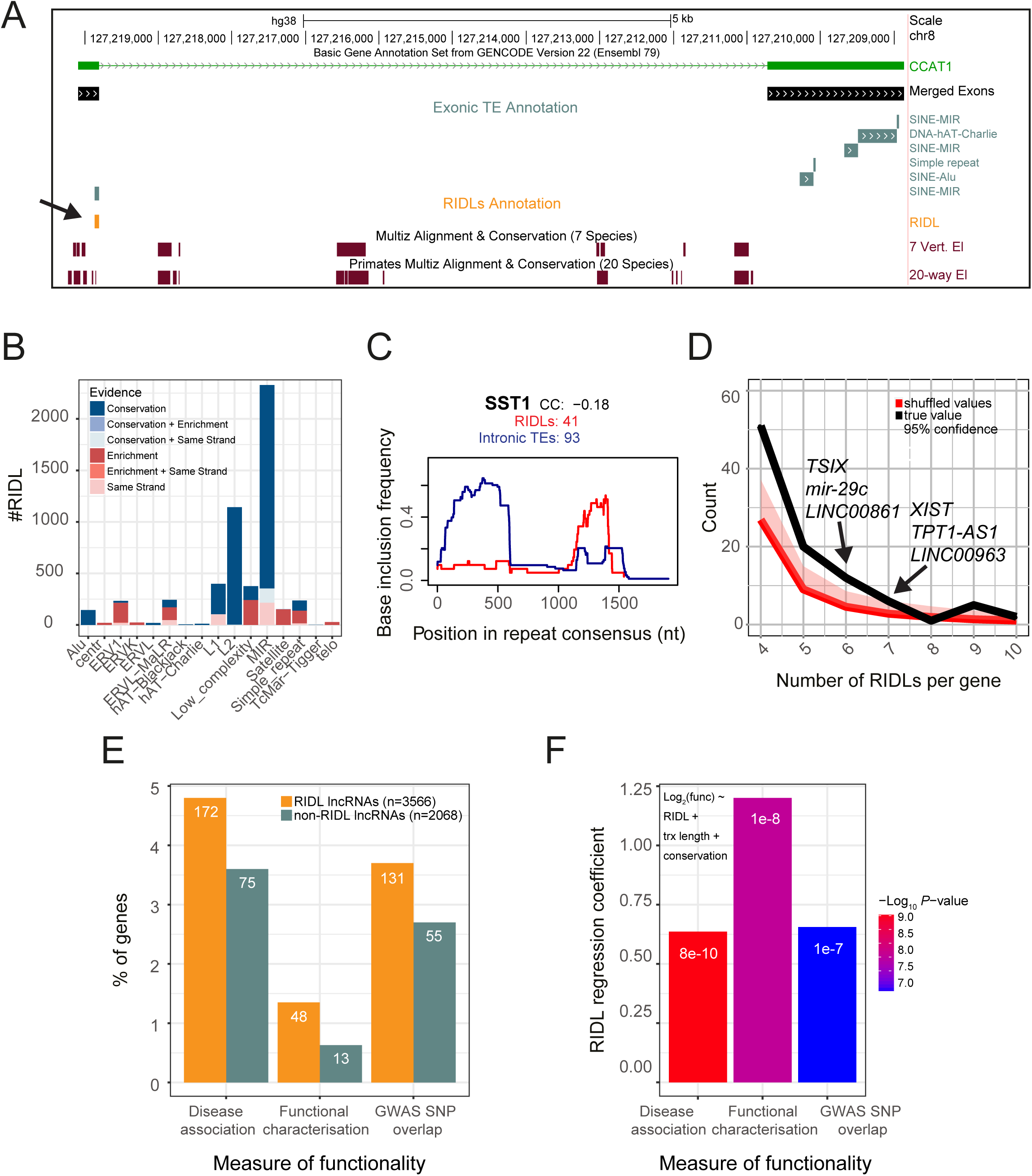
Annotated RIDLs and RIDL-lncRNAs. (A) Example of a RIDL-lncRNA gene: *CCAT1*. Of note is that although several exonic TE instances are identified (grey), including three separate MIR elements, only one is defined a RIDL (orange) due to overlap of a conserved element. (B) Breakdown of RIDL instances by TE family and evidence sources. (C) Insertion profile of SST1 RIDLs (blue) and intronic insertions (red). *x* axis shows the entire consensus sequence of SST1. *y* axis indicates the frequency with which each nucleotide position is present in the aggregate of all insertions. “CC”: Spearman correlation coefficient of the two profiles. “RIDLs” / “Intronic TEs” indicate the numbers of individual insertions considered for RIDLs / intronic insertions, respectively. (D) Number of lncRNAs (*y* axis) carrying the indicated number of RIDL (*x* axis) given the true distribution (black) and randomized distribution (red). The 95% confidence interval was computed empirically, by randomly shuffling RIDLs across the entire lncRNA annotation. (E) Percentage of RIDL-lncRNAs, and a length-matched set of non-RIDL lncRNAs, which are present in disease-and cancer-associated lncRNA databases (see Materials and Methods), in the lncRNAdb database of functional lncRNAs (36), or contain at least one trait/disease-associated SNP in an exonic region. Numbers denote gene counts. (F) Plot shows regression coefficients for the “RIDL” term in the indicated multiple logistic regression model using the same measures of functionality than in (E). Colours indicate the associated *p*-value. These values assess the correlation between RIDL number and measures of functionality of their host transcript, while accounting for transcript length (trx length) and conservation.

After removing redundancy, the final RIDL annotation consists of 5374 elements, located within 3566 distinct lncRNA genes (Figure 3D). These represent 12% (5374/46474) of all exonic TE fragments. The most predominant TE families are MIR and L2 repeats, representing 2329 and 1143 RIDLs (Figure 4B). The majority of both are defined based on evolutionary evidence (Figure 4B, Supplemental Figure S3). In contrast, RIDLs composed by ERV1, low complexity, satellite and simple repeats families are more frequently identified due to exonic enrichment (Figure 4B). The entire RIDL annotation is available in Supplemental File S1.

It is important to consider this RIDL annotation as an “enrichment group”, with a greater proportion of functional TEs than when using the entire exonic TE set. Using introns as a reference, we conservatively estimate the fraction of true positive predictions to range from 12% (strand bias) to 40% (phastCons primate) and 78% (exonic enrichment) (Supplemental Figure S4).

We also examined the evolutionary history of RIDLs. Using 6-mammal alignments, their depth of evolutionary conservation could be inferred (Supplemental Figure S5). 12% of instances appear to be Great Ape-specific, with no orthologous sequence beyond chimpanzee. 47% are primate-specific, while the remaining 40% are identified in at least one non-primate mammal. The wide timeframe for appearance of RIDLs is consistent with the wide diversity of TE types, from ancient MIR elements to presently-active LINE1 (Jurka et al. 1995; Smith et al. 2013; Konkel et al. 2010).

Instances of genomic TE insertions typically represent a fragment of the full consensus sequence. We hypothesised that particular regions of the TE consensus will be important for RIDL activity, introducing selection for these regions that would distinguish them from unselected, intronic copies. To test this, we compared insertion profiles of RIDLs to intronic instances, for each TE type, and used the correlation coefficient (CC) as a quantitative measure of similarity (Figure 4C and Supplemental File S2). For 17 cases, a CC<0.9 points to possible selective forces acting on RIDL insertions. An example is the macrosatellite SST1 repeat where RIDL copies in 41 lncRNAs show a strong preference inclusion of the 3′ end, in contrast to the general 5′ preference observed in introns (Figure 4C). This suggests a possible functional relevance for the 1000-1500 nt region of the SST1 consensus.

To assess whether RIDLs experience purifying evolutionary selection in modern humans, we analysed the derived allele frequency (DAF) spectrum of their overlapping SNPs (Supplemental Figure S6) (Haerty and Ponting 2013)(Tan et al. 2017). This showed that RIDLs (orange bars) have a greater proportion of rare (DAF<0.1) alleles compared to other TEs in exons (green bars) or introns (turquoise bars) of the same lncRNAs, and indeed compared to non-RIDL exonic nucleotides (black bars). These differences fail to reach statistical significance, possibly due to small sample sizes. Overall these data are consistent with RIDLs experiencing an elevated rate of purifying evolutionary selection in modern humans compared to nearby neutral sequence, although larger datasets will be required before this can be stated conclusively.

### RIDL-carrying lncRNAs are enriched for functions and disease roles

We next looked for evidence to support the RIDL annotation by investigating the properties of their host lncRNAs. We first asked whether RIDLs are randomly distributed amongst lncRNAs, or else non-randomly clustered in a smaller number of genes. Figure 4D shows that the latter is the case, with a significant deviation of RIDLs from a random distribution. These lncRNAs carry a mean of 1.15 RIDLs / kb of exonic sequence (median: 0.84 RIDLs/kb) (Supplemental Figure S7).

Are RIDL-lncRNAs more likely to be functional? To address this, we compared lncRNA genes carrying one or more RIDLs, to a length-matched set of control lncRNAs (Figure 4E, Supplemental Figure S8). We observed that RIDL-lncRNAs are (1) over-represented in the reference database for functional lncRNAs, lncRNAdb (Quek et al. 2015), (2) enriched in associations with cancer and other diseases, and (3) enriched in their exons for trait/disease-associated SNPs. In order to estimate the impact of carrying RIDLs on the functional-associated outcomes mentioned above, while controlling for potential biases from conservation and length, we performed multiple logistic regression analysis. In each case, the overlap with RIDL-lncRNAs was positive and statistically significant (Figure 4F). However we did not observed any difference in mean or maximum expression of RIDL-lncRNAs to length matched controls across ten tissues of the Human Body Map RNA-seq dataset (Supplemental Figure S9).

In addition to *CCAT1* (Figure 4A) (Nissan et al. 2012) there are a number of deeply-studied RIDL-containing genes. *XIST*, the X-chromosome silencing RNA contains seven internal RIDL elements. As we pointed out previously (Johnson and Guigó 2014) these include an array of four similar pairs of MIRc / L2b repeats. The prostate cancer-associated *UCA1* gene has a transcript isoform promoted from an LTR7c, as well as an additional internal RIDL, thereby making a potential link between cancer gene regulation and RIDLs. The *TUG1* gene, involved in neuronal differentiation, contains highly evolutionarily-conserved RIDLs including Charlie15k and MLT1K elements (Johnson and Guigó 2014). Other RIDL-containing lncRNAs include *MEG3, MEG9, SNHG5, ANRIL, NEAT1, CARMEN1* and *SOX2OT*. *LINC01206*, located adjacent to *SOX2OT*, also contains numerous RIDLs. A full list can be found in Supplemental File S3.

### Correlation between RIDLs and subcellular localisation of host transcript

The location of a lncRNA within the cell is of key importance to its molecular function (Derrien et al. 2012; Cabili et al. 2015)(Mas-Ponte et al. 2017), therefore we next investigated whether RIDLs might regulate lncRNA localisation (Zhang et al. 2014a; Hacisuleyman et al. 2016b)(Chillón and Pyle 2016) (Figure 5A). Using subcellular RNA-seq data based on 10 ENCODE cell lines (Djebali et al. 2012), we calculated the relative nuclear/cytoplasmic localisation in log2 units, or “Relative Concentration Index” (RCI) (Mas-Ponte et al. 2017). Using this dataset, we tested each of the 99 RIDL types for association with localisation of their host transcript.

**Figure 5:**
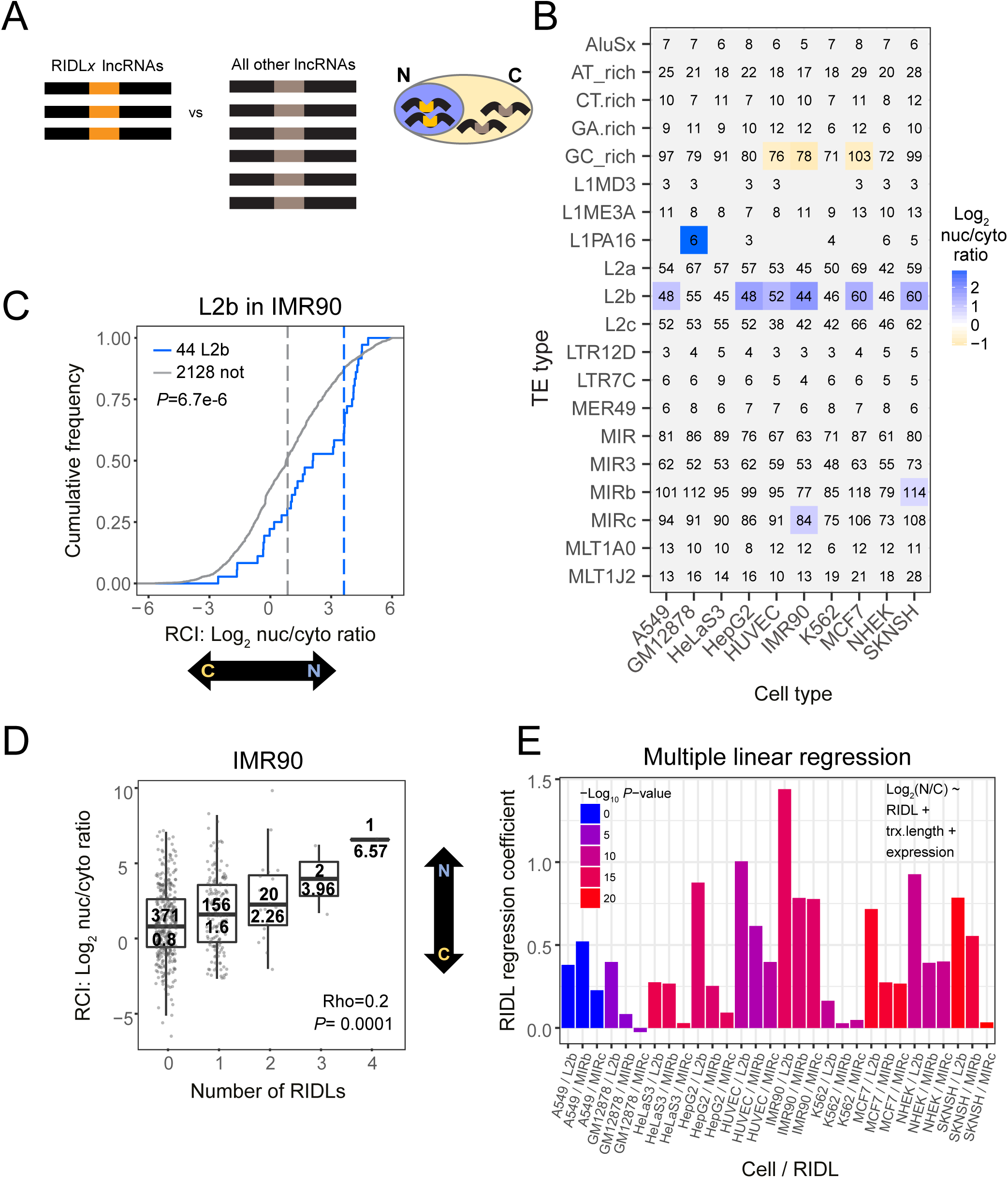
Correlation between RIDLs and host lncRNA nuclear/cytoplasmic localisation. (A) Outline of in silico screen for localisation-regulating RIDLs. For each RIDL-type / cell-type combination, the nuclear/cyto-plasmic localisation of RIDL-lncRNAs is compared to all other detected lncRNAs. (B) Results of an in silico screen. Rows: RIDL types; Columns: Cell types. Significant RIDL-cell type combinations are coloured (Benjamini-Hochberg corrected *p*-value < 0.01; Wilcoxon test). Colour scale indicates the nuclear/cytoplasmic ratio mean of RIDL-lncRNAs. Numbers in cells indicate the number of considered RIDL-lncRNAs. Analyses were performed using a single representative transcript isoform from each gene locus, being that with the greatest number of exons. (C) The nuclear/cytoplasmic localization of lncRNAs carrying L2b RIDLs in IMR90 cells. Blue indicates lncRNAs carrying ≥1 RIDLs, grey indicates all other detected lncRNAs (“not”). Dashed lines represent medians. Significance was calculated using Wilcoxon test (*P*). (D) The nuclear/cytoplasmic ratio of lncRNAs as a function of the number of RIDLs that they carry (L1PA16, L2b, MIRb, MIRc). Correlation coefficient (Rho) and corresponding *p*-value (*P*) were calculated using Spearman correlation, two-sided test. In each box, upper value indicates the number of lncRNAs, and lower value the median. (E) Plot shows regression coefficients for the “RIDL” term in the indicated linear model using L2b, MIRb and MIRc RIDLs (see Methods). Colours indicate the associated *p*-value. These values assess the correlation between RIDL number and nuclear/cytoplasmic localisation (Log2(N/C)) of their host transcript, while accounting for possible confounding factors of transcript length (trx.length) or whole-cell expression levels (expression).

After correcting for multiple hypothesis testing using the Benjamini-Hochberg method (Benjamini and Hochberg 1995), this approach identified four distinct RIDL types: L1PA16, L2b, MIRb and MIRc (Figure 5B). For example, 44 lncRNAs carrying L2b RIDLs have 6.9-fold higher relative nuclear/cytoplasmic ratio in IMR90 cells, and this tendency is observed in six different cell types (Figure 5B,C).

The degree of nuclear localisation increases in lncRNAs as a function of the number of RIDLs (L1PA16, L2b, MIRb and MIRc) they carry (Figure 5D). We also found a significant relationship between GC-rich elements and cytoplasmic enrichment across three independent cell samples. The GC-rich-containing lncRNAs have between 2 and 2.3-fold higher relative expression in the cytoplasm of these cells (Supplemental Figure S10).

We were curious whether this relationship with localisation is only a property of RIDLs, or conversely, holds true when considering any instances of L1PA16, L2b, MIRb and MIRc. Indeed when the preceding analysis was repeated with unfiltered TE instances, the latter was observed (Supplemental Figure S11). However, the strength of the effect was consistently lower than for RIDLs (Supplemental Figure S12). This difference between RIDLs and unfiltered TEs supports both the usefulness of the RIDL identification method, and the idea that RIDLs are under selection as a result of their effect on localisation.

We were concerned that two un-modelled confounding factors that positively correlated with TE number could explain the observed data: transcript length and whole-cell gene expression. To address this, we performed multiple linear regression for localisation with explanatory variables of RIDL number, transcript length and whole-cell expression (Figure 5E). Such a model accounts independently for each variable, enabling one to eliminate confounding effects. Training such models for each cell type / RIDL pair, we observed positive and statistically-significant contributions for RIDL number in most cases. We also observed weaker but significant contributions from transcript length and whole-cell expression terms, indicating that our intuition was correct that these factors influence localisation independent of RIDLs (Supplemental Figure S13A,B). We drew similar conclusions from equivalent analyses using partial correlation analysis (Supplemental Figure S13C). In summary, observed RIDLs correlate with lncRNA localisation even when controlling for other factors,

Given that L2b and MIR elements predate human-mouse divergence, we attempted to perform similar analyses in mouse cells. However given that just two equivalent datasets are available at present (Bahar Halpern et al. 2015)(Tan et al. 2015), as well as the relatively low number of annotated lncRNAs in mouse, we were unable to draw statistically-robust conclusions regarding the evolutionary conservation of this phenomenon.

### Intra-gene correlation between RIDLs and subcellular localisation

LncRNA gene loci are often composed of multiple, differentially-spliced transcript isoforms that partially differ in their mature sequence. We reasoned that differential inclusion of RIDL-containing exons should give rise to differences in localisation amongst transcripts from the same gene locus. In other words, for RIDL-lncRNA gene loci having multiple transcript isoforms, those isoforms *with* a RIDL should display greater nuclear enrichment than those isoforms *without* a RIDL (Figure 6, left panel).

**Figure 6:**
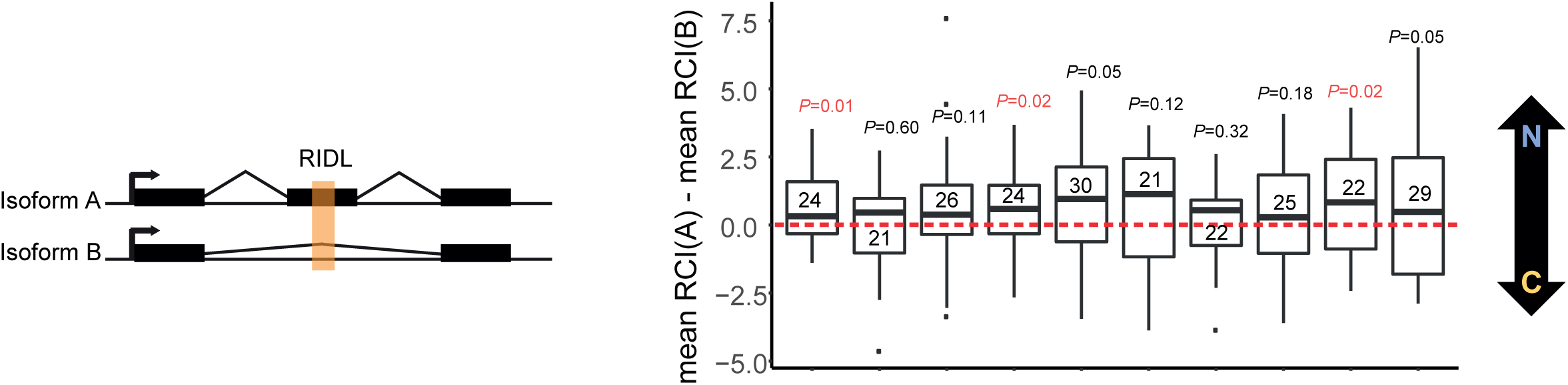
RIDLs correlate with differential localisation of lncRNA transcripts from the same locus. Distribution of differences between RCI mean of transcripts with nuclear RIDL (mean RCI(A)) and RCI mean of transcripts without nuclear RIDL (mean RCI(B)). A positive value indicates that RIDL-carrying transcripts are more nuclear-enriched than non-RIDL transcripts. Data were calculated individually for every gene that has ≥1 RIDL-transcript and ≥1 non-RIDL transcript expressed in a given cell line. Numbers inside the boxplots indicate the number of gene loci analysed for each cell line. Horizontal bar indicates the median. Here “nuclear RIDL” refers to L1AP16, MIRb, MIRc and L2b. *P*-values obtained from one-sided t-test are shown (in red when *P <* 0.05).

We tested this individually for each cell type. For every appropriate RIDL-lncRNA locus (numbers shown inside boxplot), we calculated the difference in the mean of the localisation between their RIDL and non-RIDL isoforms (Figure 6, right panel). For every cell line, the median difference was positive, indicating that RIDL-carrying transcript isoforms are more nuclear enriched than their non-RIDL cousins from the same gene locus. Given our *a priori* hypothesis that RIDLs promote nuclear enrichment, statistical significance was tested by comparison to zero using a 1-sided *t*-test. Altogether these data point to a consistent correlation between the presence of certain exonic TE elements, L1PA16, L2b, MIRb and MIRc, and the nuclear enrichment of their host lncRNA.

### RIDLs play a causative role in lncRNA nuclear localisation

To more directly test whether RIDLs play a causative role in nuclear localisation, we designed an experimental approach to quantify the effect of exonic TEs on localisation of a transfected lncRNA. We selected three lncRNAs, based on: (i) presence of L2b, MIRb and MIRc RIDLs; (ii) moderate expression; (iii) nuclear localisation, as inferred from RNA-seq (Figure 7A,B and Supplemental Figure S14). Nuclear localisation of these candidates could be validated in HeLa cells using qRTPCR (Figure 7C).

**Figure 7:**
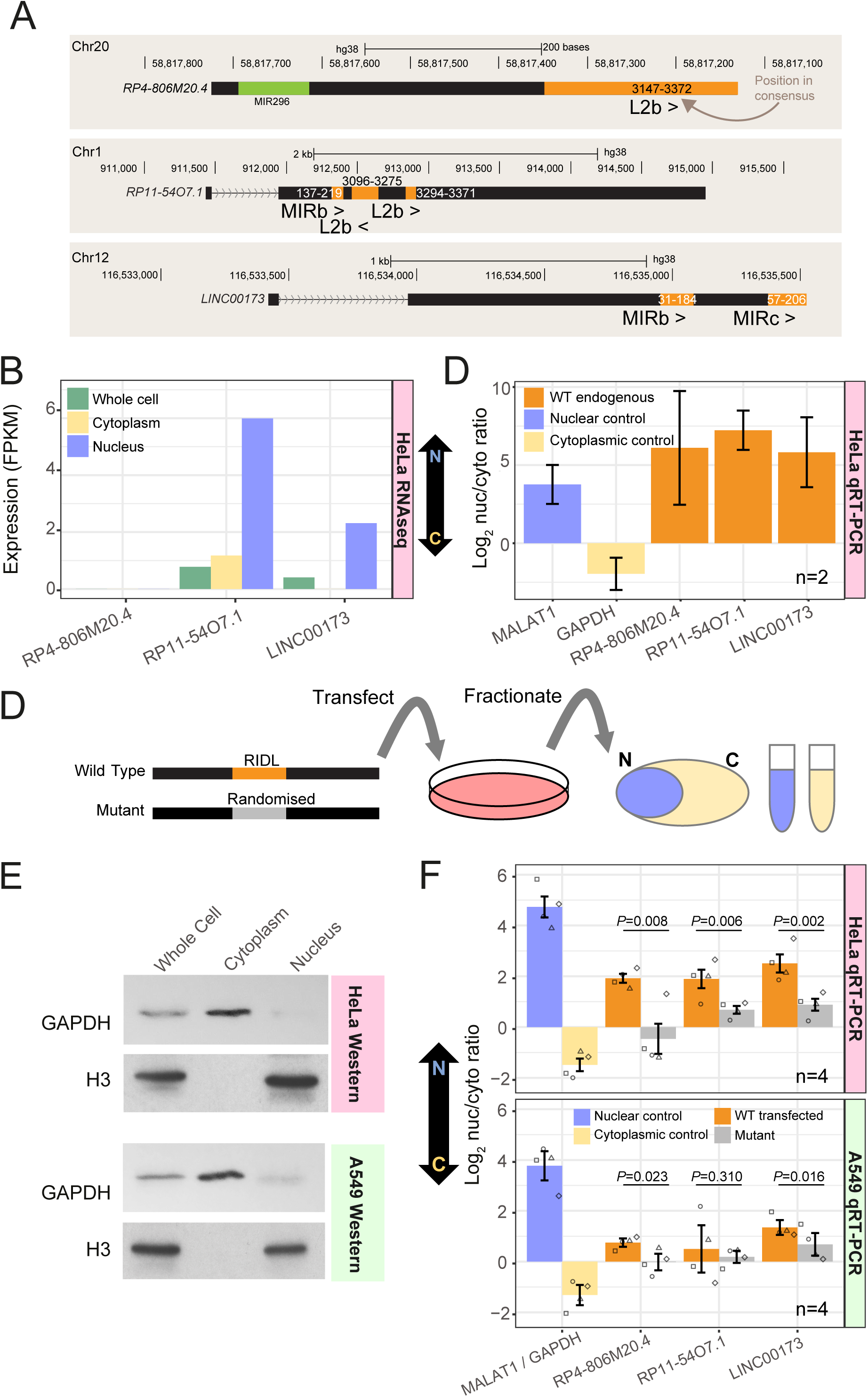
Disruption of RIDLs results in lncRNA relocalisation from nucleus to cytoplasm. (A) Structures of candidate RIDL-lncRNAs. Orange indicates RIDL positions. For each RIDL, numbers indicate the position within the TE consensus, and its orientation with respect to the lncRNA is indicated by arrows (“>“ for same strand, “<“ for opposite strand). (B) Expression of the three lncRNA candidates as inferred from HeLa RNAseq (40). (C)Nuclear/cytoplasmic localisation of endogenous candidate lncRNA copies in wild-type HeLa cells, as measured by qRT-PCR. (D) Experimental design. (E) The purity of HeLa and A549 subcellular fractions was assessed by Western blotting against specific markers. GAPDH / Histone H3 proteins are used as cytoplasmic / nuclear markers, respectively. (F) Nuclear/cytoplasmic localisation of transfected candidate lncRNAs in HeLa (upper panel) and A549 (lower panel). GAPDH/MALAT1 are used as cytoplasmic/nuclear controls, respectively. N indicates the number of biological replicates (values from all replicates are plotted, each replicate is represented by a different dot shape), and error bars represent standard error of the mean. *P*-values for paired t-test (1 tail) are shown.

We formulated an assay to compare the localisation of transfected lncRNAs carrying wild-type RIDLs, and mutated versions where the RIDL sequence was randomised without altering sequence composition (“Mutant”) (Figure 7D, full sequences available in Supplemental File S4). Wild-type and Mutant lncRNAs were transfected into cultured cells and their localisation evaluated by fractionation. qRT-PCR primers were designed to distinguish transfected Wild-Type and Mutant transcripts from endogenously-expressed copies. Transgenes were typically expressed in a range of 0.2-to 10-fold compared to their endogenous transcripts (Supplemental Figure S15). Fractionation purity was verified by Western blotting (Figure 7E) and qRT-PCR (Figure 7F), and stringent DNase-treatment ensured that plasmid DNA made negligible contributions to our results (Supplemental Figure S16).

With this setup, we compared the nuclear/cytoplasmic localisation of lncRNAs with and without exonic RIDL sequences (Figure 7F). We observed a potent and consistent impact of RIDLs on nuclear/cytoplasmic localisation in HeLa cells: for all three candidates, loss of RIDL sequence resulted in relocalisation of the host transcript from nucleus to cytoplasm (Figure 7F, upper panel). We repeated these experiments in another cell line, A549, and observed similar, albeit less pronounced, effects (Figure 7F, lower panel). This difference may be due to the less nuclear localisation of the endogenous transcripts in A549 (Supplemental Figure S17). To summarise, exonic L2b, MIRb and MIRc elements promote the nuclear enrichment of host lncRNAs.

## Discussion

Recent years have seen a rapid increase in the number of annotated lncRNAs. However, our understanding of their molecular functions, and how such functions are encoded in primary RNA sequences, lag far behind. Two recent conceptual developments offer hope for resolving the sequence-function code of lncRNAs: First, the idea that the subcellular localisation of lncRNAs is a readily quantifiable characteristic that holds important clues to function; Second, that the abundant transposable element (TE) content of lncRNAs may contribute to functionality.

In this study, we have linked these two ideas, by showing evidence that certain TEs can drive the nuclear enrichment of lncRNAs. A global correlation analysis of TEs and RNA localisation data revealed a handful of TEs, most notably LINE2b, MIRb and MIRc, which positively and significantly correlate with the degree of nuclear/cytoplasmic localisation of their host transcripts. This correlation is observed in multiple cell types, and scales with the number of TEs present. A causative link was established experimentally, confirming that the indicated TEs are sufficient for a two-to four-fold increase in nuclear/cytoplasmic localisation. There are two principal explanations for this phenomenon. First, an “active” process whereby TEs are recognised by a cellular transport pathway, as demonstrated for *Alus* by Lubelsky and Ulitsky (Lubelsky and Ulitsky 2018). Second, a “passive” process where TEs destabilise transcripts leading to a concentration gradient from nucleus to cytoplasm. Although future studies will examine this question in detail, the fact that we do not observe a constant difference in steady-state levels in TE/mutated transgenes, would be more consistent with the active model.

These data support the hypothesis that exonic TE elements can act as functional lncRNA domains. In this “RIDL hypothesis”, transposable elements are co-opted by natural selection to form “Repeat Insertion Domains of LncRNA”, that is, fragments of sequence that confer adaptive advantage through some change in the activity of their host lncRNA. We proposed that RIDLs may serve as binding sites for proteins or other nucleic acids, and indeed a growing body of evidence supports this (reviewed in (Johnson and Guigó 2014)). In the context of localisation, RIDLs could mediate nuclear retention through hybridisation to complementary repeats in genomic DNA or through their described interactions with nuclear proteins (Kelley et al. 2014). In the course of this study we bioinformatically identified five candidate proteins (HNRNPU, HNRNPH2, HuR, KHDRBS1, TARDBP), however we could not find evidence that they contribute to RIDL-lncRNA localisation. Identification of any proteins that mediate RIDLs’ localisation activity may be achieved in future through pulldown approaches (Marín-Béjar and Huarte 2015).

The localisation RIDLs discovered – MIR and LINE2-are both ancient and contemporaneous, being active prior to the mammalian radiation (Cordaux and Batzer 2009). Both have previously been associated with acquired roles in the context of genomic DNA, but not to our knowledge in RNA (Jjingo et al. 2014)(Johnson et al. 2006). Although the evolutionary history of lncRNAs remains an active area of research and accurate dating of lncRNA gene birth is challenging, it appears that the majority of human lncRNAs were born after the mammalian radiation (Hezroni et al. 2015)(Hezroni et al. 2017)(Necsulea et al. 2014)(Washietl et al. 2014). This would mean that MIR and LINE2 RIDLs were pre-existing sequences that were exapted by newly-born lncRNAs, corresponding to the “latent” exaptation model proposed by Feschotte and colleagues (Chuong et al. 2017). However it is also possible that for other cases the reverse could be true – a pre-existing lncRNA exapts a newly-inserted TE. Given that nuclear retention is at odds with the primary needs of natural TE transcripts to be exported to the cytoplasm, we propose that the observed nuclear localisation activity is a more modern feature of L2b/MIR RIDLs, which is unrelated to their original roles.

Our approach for identifying localisation-regulating RIDLs has advantages over previous studies (Lubelsky and Ulitsky 2018; Hacisuleyman et al. 2016c) in terms of its genome-wide scale. However an unavoidable consequence of our use of evolutionary conservation as a filter, is that it likely biases our analysis against recently-evolved TEs such as *Alus*. It remains entirely possible that modern TEs also influence lncRNA localisation, but cannot be detected using the signals of selection that we have employed. On the other hand, MIRb and MIRc were only identified in one cell type each. We expect this reflects low sensitivity of the statistical screen, rather than cell-type specificity alone, because (i) in a focussed re-analysis (Supplemental Figure S12) the effect was observed in multiple cells, and (ii) experimental validation confirmed it in two independent cell types (Figure 7F).

This is further supported by the recent study of Lubelsky and Ulitsky, who performed an experimental screen for localisation motifs in 37 nuclear-enriched lncRNAs, and identified *Alu*Sx as a nuclear-localisation element (Lubelsky and Ulitsky 2018). These 37 lncRNAs are enriched for RIDLs (62% of Lubelsky lncRNAs contain at least one RIDL, compared to 22% for other Gencode v21 lncRNAs, P=4e-6, Fisher exact test), as well as for the three localisation RIDLs identified here (L2b, MIRb, MIRc: 32% vs 9%, P=3e-4) (Supplemental Figure S18A). Although our bioinformatic screen did not identify *Alu*Sx, a naive unfiltered re-analysis of our data supports Lubelsky’s experimental finding that *Alu*Sx-carrying lncRNAs tend to be more nuclear across multiple cell types (Supplemental Figure S18B). Together, these considerations open the possibility that other localisation-controlling TE types may await discovery.

More generally, the RIDL predictions showed rather low concordance between the various selection evidence used (Figure 3D). This likely reflects a number of factors: young evolutionary age of some of the most common TEs, generally low statistical power due to large background of neutral TEs and multiple hypothesis testing, and false positives due to TEs that promote transcription or splicing of lncRNAs. However it is worthy of note that validated candidates L2b, MIRb and MIRc are all implicated by multiple, independent evidence sources (Figure 3D).

This work marks a step in the ongoing efforts to map the domains of lncRNAs. Previous studies have utilised a variety of approaches, from integrating experimental protein-binding data (Van Nostrand et al. 2016)(Hu et al. 2017)(Li et al. 2014), to evolutionarily-conserved segments (Smith et al. 2013)(Seemann et al. 2017). Previous maps of TEs have highlighted their profound roles in lncRNA gene evolution (Kapusta et al. 2013)(Kelley and Rinn 2012)(Hezroni et al. 2015). However, the present RIDL annotation stands apart in attempting to identify the subset of TEs with evidence for selection. We hope that this RIDL map will prove a resource for future studies to better understand functional domains of lncRNAs. Although various evidence suggests that the RIDL annotation is a useful enrichment group of functional TE elements, it contains a substantial false positive (and likely also false negative) rates that will have to be improved in future.

This study may help to explain a longstanding and unexplained property of lncRNAs: their nuclear enrichment (Derrien et al. 2012)(Ulitsky and Bartel 2013). Although they are readily detected in the cytoplasm, lncRNAs general tendency is to have higher nuclear/cytoplasmic ratios compared to mRNAs (Clark et al. 2012)(Ulitsky and Bartel 2013)(Derrien et al. 2012)(Mas-Ponte et al. 2017). This is true across various human and mouse cell types. Although this may partially be explained by decreased stability (Mukherjee et al. 2017), it is likely that RNA sequence motifs also contribute to nuclear localisation (Chillón and Pyle 2016)(Zhang et al. 2014a). Here we show that this is the case, and that the enrichment of certain RIDL types in lncRNA mature sequences is likely to be a major contributor to lncRNA nuclear retention. In contrast, the far lower exonic content of TEs in protein-coding mRNAs may help explain their greater cytoplasmic abundance (Kapusta and Feschotte 2014). Indeed, even within the cytoplasm, there is evidence that TE content may also influence the efficiency with which lncRNAs are trafficked to the translation machinery (Carlevaro-Fita et al. 2016). Together, this evidence may reflect unknown cellular quality control mechanisms that vet RNAs based on their TE content, tending to retain TE-rich sequences (including lncRNAs or incorrectly processed mRNAs) in the nucleus, and promote the cytoplasmic export and ribosomal loading of canonical TE-poor mRNAs.

In summary therefore, we have made available a first annotation of selected RIDLs in lncRNAs, and described a new paradigm for TE-derived fragments as drivers of nuclear localisation in lncRNAs.

## Materials and Methods

All operations were carried out on human genome version GRCh38/hg38, unless stated otherwise.

### Exonic TE curation

*RepeatMasker* annotations were downloaded from the UCSC Genome Browser (version hg38) on December 31st 2014, and GENCODE version 21 lncRNA annotations in GTF format were downloaded from www.gencodegenes.org (Harrow et al. 2012). Annotations were not filtered further. The ‘*transposon.profiler’* script, largely based on BEDTools’ *intersect* and *merge* functionalities (Quinlan and Hall 2010), was used to annotate exonic and intronic TEs of the given gene annotation (Supplemental File S5). Exons of all transcripts belonging to the given gene annotation were merged, henceforth referred to as “exons”. The set of introns was curated by subtracting the merged exonic sequences from the full gene spans, and only retaining those introns that belonged to a single gene. Intronic regions were assigned the strand of the host gene.

The *RepeatMasker* annotation file was intersected with exons and classified into one of 6 categories: TSS (transcription start site), overlapping the first exonic nucleotide of the first exon; splice acceptor, overlapping exon 5′ end; splice donor, overlapping exon 3′ end; internal, residing within an exon and not overlapping any intronic sequence; encompassing, where an entire exon lies within the TE; TTS (transcription termination site), overlapping the last nucleotide of the last exon. In every case, the TEs are separated by strand relative to the host gene: + where both gene and TE are annotated on the same strand, otherwise -. The result is the “Exonic TE Annotation” (Supplemental File S6).

### RIDL identification

Using this Exonic TE Annotation, we identified the subset of individual TEs with evidence for functionality. For certain analysis, an Intronic TE Annotation was also employed, being the output for the equivalent intron annotation described above. Three different types of evidence were used: enrichment, strand bias and evolutionary conservation.

In enrichment analysis, the exon/intron ratio of the fraction of nucleotide coverage by each repeat type was calculated. Any repeat type with >2-fold exon/intron ratio was considered as a candidate. All exonic TE instances belonging to such TE types are defined as RIDLs.

In strand bias analysis, a subset of Exonic TE Annotation was used, being the set of non-splice junction crossing TE instances (“noSJ”). This additional filter was employed to guard against false positive enrichments for TEs known to provide splice sites (Sela et al. 2007; Lev-Maor et al. 2003). For all TE instances, the “relative strand” was calculated: positive, if the annotated TE strand matches that of the host transcript; negative, if not. Then for every TE type, the ratio of relative strand sense/antisense was calculated. Statistical significance was calculated empirically: entire gene structures were randomly re-positioned in the genome using *BEDTools shuffle*, and the intersection with the entire *RepeatMasker* annotation was re-calculated. For each iteration, sense/antisense ratios were calculated for all TE types. A TE type was considered to have significant strand bias, if its true ratio exceeded (positively) all of 1000 simulations. All exonic instances of these TE types that also have the same strand orientation to the host transcript are defined as RIDLs. On the other hand, after inspection of the data, we decided to exclude TEs with significant antisense enrichment. This is because most instances were from the LINE1 class, which are known to interfere with gene expression when falling on the same strand (Perepelitsa-Belancio and Deininger 2003). Therefore, we considered it likely that observed antisense enrichment is simply an artefact of selection against insertion on the same strand, and in the interests of controlling the false positive prediction rate, decided to exclude these cases.

In evolutionary analysis, four different annotations of evolutionarily-conserved regions were treated similarly, using unfiltered Exonic TE Annotations. Primate, Placental Mammal and Vertebrate phastCons elements based on 46-way alignments were downloaded as BED files from UCSC Genome Browser (Siepel et al. 2005), while the ECS conserved regions from obtained from Supplemental Data of Smith et al (Smith et al. 2013) (see Supplemental File S7 for summary). Because at the time of analysis, phastCons elements were only available for hg19 genome build, we mapped them to hg38 using *LiftOver* utility. For each TE type we calculated the exonic/intronic conservation ratio. To do this we used *IntersectBED* (Quinlan and Hall 2010) to overlap exonic locations with TEs, and calculate the total number of nucleotides overlapping. We performed a similar operation for intronic regions. Then for each TE type, we calculated the ratio of conserved TE nucleotides for exons compared to introns:

Relative exonic-intronic conservation (REIC) = (*C*_*e*_ / (*C*_*e*_ + *N*_*e*_)) / (*C*_*i*_ / (*C*_*i*_ + *N*_*i*_))

Where *C* is conserved TE nucleotides, *N* is non-conserved TE nucleotides, and subscripts *e* and *i* denote exonic and intronic, respectively. Note that, because it calculates fractional overlap of TEs by conserved elements, REIC normalises for different lengths of exons and introns (Supplemental Figure S19).

To estimate the background, the conserved element BED files were positionally randomized 1000 times using *BEDTools shuffle*, each time recalculating REIC. We considered to be significantly conserved those TE types where the true REIC was greater or less than every one of 1000 randomised REIC values. All exonic instances of these TE types that also intersect the appropriate evolutionarily conserved element are defined as RIDLs. This approach of shuffling conserved elements displayed no apparent bias in the length of TEs it identifies (Supplemental Figure S2D). We also tested an alternative approach for estimating significance whereby conserved elements were held constant, and TEs were positionally randomised. While there was a significant overlap in identified candidate RIDLs, this method displayed a bias towards longer TEs (Supplemental Figure S2D), and therefore was not employed further.

We chose to randomise conserved elements, rather than TEs because the former are enriched in lncRNA exons (Pegueroles and Gabaldón 2016). Thus, using randomised TEs to estimate background REIC would lead to overestimation of exonic TE conservation, and hence underestimate the rate of conservation of TEs in real data.

All RIDL predictions were then merged using *mergeBED* and any instances with length <10 nt were discarded. The outcome, a BED format file with coordinates for hg38, is found in Supplemental File S1.

False discovery rates (FDR) were estimated for RIDL predictions. TE type FDR estimates were based on shuffling simulations described above. Empirical *p*-values for true data were estimated according to *P*=(rank in distribution)/(1 + number of simulations). For significant cases, where the true value exceeded all n=1000 simulations, this value was conservatively defined to be *P*=0.001. These empirical *p*-values were then converted to FDR using the r command “p.adjust” with “fdr” setting. Accordingly, empirical significance cutoff (*P*<0.001) mentioned in the main text corresponds to the following FDR values: Strand bias: 0.027; Vertebrate phastCons: 0.013; Placental phastCons: 0.014; Primate phastCons: 0.009; ECS: This analysis is conservative, since empirical *p*-values of candidates are rounded up in every case to 0.001.

FDR rates were also estimated at the element level. Here, the set of significant TEs were grouped for each evidence type. Then, the frequency of overlap of these TEs with the evidence type was compared for lncRNA exons and introns. This data is shown in Supplemental Figure S4.

### RIDL orthology analysis

In order to assess evolutionary history of RIDLs, we used chained alignments of human to chimp (hg19ToPanTro4), macaque (hg19ToRheMac3), mouse (hg19ToMm10), rat (hg19ToRn5), and cow (hg19ToBosTau7). Due to availability of chain files, RIDL coordinates were first converted from hg38 to hg19. Orthology was defined by *LiftOver* utility used at default settings.

### Derived allele frequency (DAF) analysis

We used allele frequencies from African population provided by the 1000 Genomes Project (Auton et al. 2015), as performed previously by (Haerty and Ponting 2013). DAF was determined for human common SNPs from dbSNP (build 150) (Sherry et al. 2001) for every group analysed. Ancestral repeats (AR) were defined as human repeats (excluding simple repeats) intersecting at least 1 nucleotide of mouse repeats defined by LiftOver, and falling within 5kb of but not overlapping RIDL-containing genes.

### Comparing RIDL-carrying lncRNAs versus other lncRNAs

In order to test for functional enrichment amongst lncRNAs hosting RIDLs, we tested for statistical enrichment of the following traits in RIDL-carrying lncRNAs compared to other lncRNAs (see below) by Fisher’s exact test:

A) Functionally-characterised lncRNAs: lncRNAs from GENCODE v21 that are present in lncRNAdb (Quek et al. 2015).

B) Disease-associated genes: lncRNAs from GENCODE v21 that are present in at least in one of the following databases or public sets: LncRNADisease (Chen et al. 2013), Lnc2Cancer (Ning et al. 2016), Cancer LncRNA Census (CLC) (Carlevaro-Fita et al. bioRxiv doi: 10.1101/152769)

C) GWAS SNPs: We collected SNPs from the NHGRI-EBI Catalog of published genome-wide association studies (Welter et al. 2014; Hindorff et al. 2009) (https://www.ebi.ac.uk/gwas/home). We intersected its coordinates with lncRNA exons coordinates.

For defining a comparable set of “other lncRNAs” we sampled from the rest of GENCODE v21 a set of lncRNAs matching RIDL-lncRNAs’ exonic length distribution (Supplemental Figure S8). We performed sampling using *matchDistribution* script: https://github.com/julienlag/matchDistribution. In order to simultaneously control for both conservation and length, we performed multiple logistic regression analysis using glm R package, with the following structure:

Functional-association outcome ∼ RIDLs + transcript length + exonic conservation

Where functional-association outcome indicates A, B and C traits defined above; RIDLs indicates the number of RIDL instances in the host gene; transcript length indicates the projected exonic length; conservation indicates the percent of exonic lncRNA nucleotides overlapping the union of primate, placental mammal and vertebrate phastCons elements. We did not find evidence for multicollinearity in any case (variance inflation factors (VIF) <1.1).

### Subcellular localisation analysis

Processed RNA-seq data from human cell fractions were obtained from ENCODE in the form of RPKM (reads per kilobase per million mapped reads) quantified against the GENCODE v19 annotation (Djebali et al. 2012; Mas-Ponte et al. 2017). Only transcripts common to both the v21 and v19 annotations were considered. For the following analysis only one transcript per gene was considered, defined as the one with largest number of exons. Nuclear/cytoplasmic ratio expression for each transcript was defined as (nuclear polyA+ RPKM)/(cytoplasmic polyA+ RPKM), and only transcripts having non-zero values (at Irreproducible Discovery Rate (IDR) between samples < 1) in both were considered. These ratios were log2-transformed, to yield the Relative Concentration Index (RCI) (Mas-Ponte et al. 2017). For each RIDL type and cell type in turn, the nuclear/cytoplasmic ratio distribution of RIDL-containing to non-RIDL-containing lncRNAs was compared using Wilcoxon test. Only RIDLs having at least three expressed transcripts in at least one cell type were tested. Resulting *p*-values were globally adjusted to False Discovery Rate using the Benjamini-Hochberg method (Benjamini and Hochberg 1995).

### Multiple linear regression and partial correlation analysis

Linear models were created in R using the “lm” package, at the level of lncRNA transcripts with the form:

localisation ∼ RIDL + transcript length + expression

Localisation refers to nuclear/cytoplasmic RCI; RIDL denotes the number of instances of a given RIDL in a transcript; expression denotes the whole cell expression level, as inferred from RNA-seq in units of RPKM. Equivalent partial correlation analyses were performed, using the R “pcor.test” function from the ‘ppcor’ package (Spearman correlation), correlating RCI with RIDL number, while controlling for transcript length and expression. We checked all regression models for multicollinearity by searching for variance inflation factors (VIF) using the ‘VIF’ command from the R package ‘fmsb’. In no case did VIF exceed 1.1, below values raising concern of multicollinearity (>4).

### Cell lines and reagents

Human cervical cancer cell line HeLa and human lung cancer cell line A549 were cultured in Dulbecco’s Modified Eagle’s medium (Sigma-Aldrich, # D5671) supplemented with 10% FBS and 1% penicillin streptomycin at 37ºC and 5 % CO2. Anti-GAPDH antibody (Sigma-Aldrich, # G9545) and anti-histone H3 antibody (Abcam, # ab24834) were used for Western blot analysis.

### Gene synthesis and cloning of lncRNAs

The three lncRNA sequences (RP11-5407, LINC00173, RP4-806M20.4) containing wild-type RIDLs, and corresponding mutated versions where RIDL sequence has been randomised (“Mutant”), were synthesized commercially (BioCat GmbH). For each gene locus, only one transcript contained the RIDL(s), and was chosen for experimental study. The sequences were cloned into pcDNA 3.1 (+) vector within the *Nhe*I and *Xho*I restriction enzyme sites. The clones were checked by restriction digestion and Sanger sequencing. The sequence of the wild-type and mutant clones are provided in Supplemental File S4.

### LncRNA transfection and sub-cellular fractionation

Wild-type and mutant lncRNA clones for each tested gene were transfected independently in separate wells of a 6-well plate. Transfections and subsequent analysis were repeated as biological replicates (four for HeLa, four for A549), defined as transfections performed on different days with different cell passages. Transfections were carried out with 2 μg of total plasmid DNA in each well using Lipofectamine 2000. 48 h post-transfection, cells from each well were harvested, pooled and re-seeded into a 10 cm dish and allowed to grow till 100% confluence. Expression of transgenes was check by qRT-PCR using specific primers, and found to typically be several-fold greater than endogenous copies (HeLa) or from 0.2-to 1-fold (A549) (Supplemental Figure S15).

The nuclear and cytoplasmic fractionation was carried out as described previously (Suzuki et al. 2010) with minor modifications. In brief, cells from 10 cm dishes were harvested by scraping and washed with 1x ice-cold PBS. For fractionation, cell pellet was re-suspended in 900 μl of ice-cold 0.1% NP40 in PBS and triturated 7 times using a p1000 micropipette. 300 μl of the cell lysate was saved as the whole cell lysate. The remaining 600 μl of the cell lysate was centrifuged for 30 sec on a table top centrifuge and the supernatant was collected as “cytoplasmic fraction”. 300 μl from the cytoplasmic supernatant was kept for RNA isolation and the remaining 300 μl was saved for protein analysis by western blot. The pellet containing the intact nuclei was washed with 1 ml of 0.1% NP40 in PBS. The nuclear pellet was re-suspended in 200 μl 1X PBS and subjected to a quick sonication of 3 pulses with 2 sec ON-2 sec OFF to lyse the nuclei and prepare the “nuclear fraction”. 100 μl of nuclear fraction was saved for RNA isolation and the remaining 100 μl was kept for western blot.

### RNA isolation and real time PCR

The RNA from each nuclear and cytoplasmic fraction was isolated using Quick-RNA MiniPrep kit (ZYMO Research, # R1055). The RNAs were subjected to on-column DNAse I treatment and clean up using the manufacturer’s protocol. For A549 samples, additional units of DNase were employed, due to residual signal in –RT samples. The RNA from each fraction was converted to cDNA using GoScript reverse transcriptase (Promega, # A5001) and random hexamer primers. The expression of each of the individual transcripts was quantified by qRT-PCR (Applied Biosystems(r) 7500 Real-Time) using indicated primers (Supplemental File S8) and GoTaq qPCR master mix (Promega, # A6001). In order to distinguish expression of transfected wild-type genes from endogenous copies, we designed forward primers against a transcribed region of the expression vector backbone. Human *GAPDH* mRNA and *MALAT1* lncRNA were used as cytoplasmic and nuclear markers, respectively. The absence of contaminating plasmid DNA in cDNA was checked for all samples using qPCR (see Supplemental Figure S16 for a representative example).

### Western Blotting

The protein concentration of each of the fractions was determined, and equal amounts of protein (50 μg) from whole cell lysate, cytoplasmic fraction, and nuclear fraction were resolved on 12 % Tris-glycine SDS-polyacrylamide gels and transferred onto polyvinylidene fluoride (PVDF) membranes (VWR, # 1060029). Membranes were blocked with 5% skimmed milk and incubated overnight at 4ºC with anti-GAPDH antibody as a cytoplasmic marker and anti p-histone H3 antibody as nuclear marker. Membranes were washed with PBS-T (1X PBS with 0.1 % Tween 20) followed by incubation with HRP-conjugated anti-rabbit or anti-mouse secondary antibodies respectively. The bands were detected using SuperSignal(tm) West Pico chemiluminescent substrate (Thermo Fisher Scientific, # 34077).

### Software availability

“*transposon.profiler*”, is available on Github at https://github.com/gold-lab/shared_scripts and in Supplemental File S5.

## Supporting information

## Acknowledgements

We wish to thank Roderic Guigó (CRG), Marc Friedlaender (SciLife Lab) and Marta Melé (Harvard) for many helpful discussions. Roberta Esposito (DBMR) and Samir Ounzain (CHUV) contributed valuable suggestions regarding experimental design and analysis. Julien Lagarde (CRG) kindly provided help in gene sampling analysis. Carlos Pulido (DBMR) assisted with RNA-seq analysis, and Reza Sodaie (CRG) helped with combinatorial analysis of TEs. We acknowledge Deborah Re (DBMR), Silvia Roesselet (DBMR) and Marianne Zahn (Inselspital) for administrative support. CN is supported by grants TIN-2013-41990-R and DPI-2017-84439-R from the Spanish Ministry of Economy, Industry and Competitiveness (MINECO). This research was funded by the NCCR “RNA & Disease” funded by the Swiss National Science Foundation, and by the Medical Faculty of the University and University Hospital of Bern.

