## Supplementary material for "Ancient exapted transposable elements promote nuclear enrichment of human long noncoding RNAs"

Supplemental Figure S1

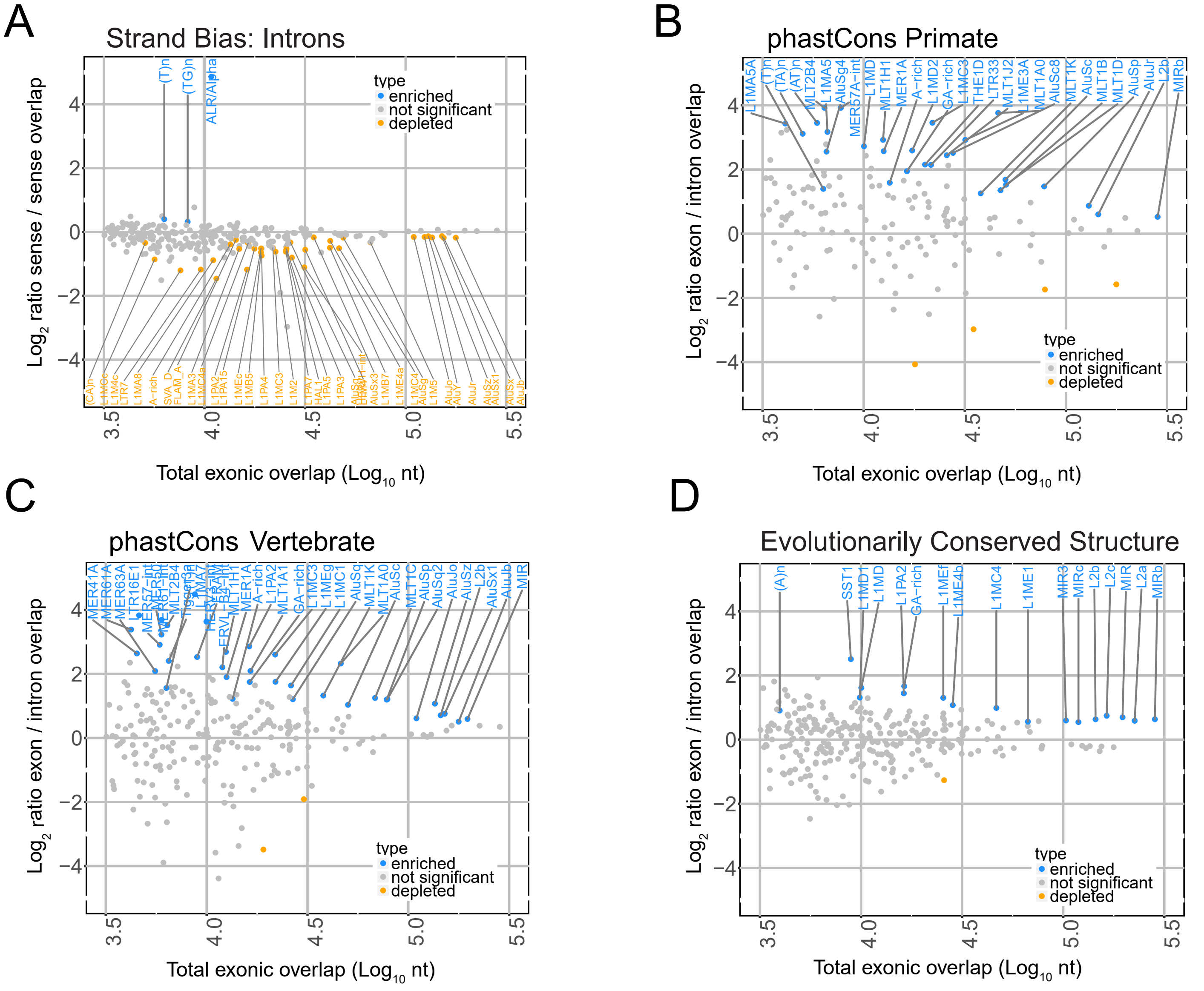


**Supplemental Figure S1:** (A) Figure shows, for every TE type, the ratio of intronic nucleotide coverage in sense vs antisense configuration. “Sense” here is defined as sense of TE annotation relative to the sense of the overlapping intron. Significantly-enriched TE types are shown in blue. Statistical significance was estimated by a randomisation procedure, and significance is defined at an uncorrected empirical p-value < 0.001 (See Material and Methods). (B-D) As for (A), but y axis records the ratio of nucleotide overlap in exons vs introns by phastCons primate-conserved elements, phastCons vertebrate-conserved elements and Evolutionarily Conserved Structures (ECS), respectively.
