## Supplementary material for "Ancient exapted transposable elements promote nuclear enrichment of human long noncoding RNAs"

Supplemental Figure S2

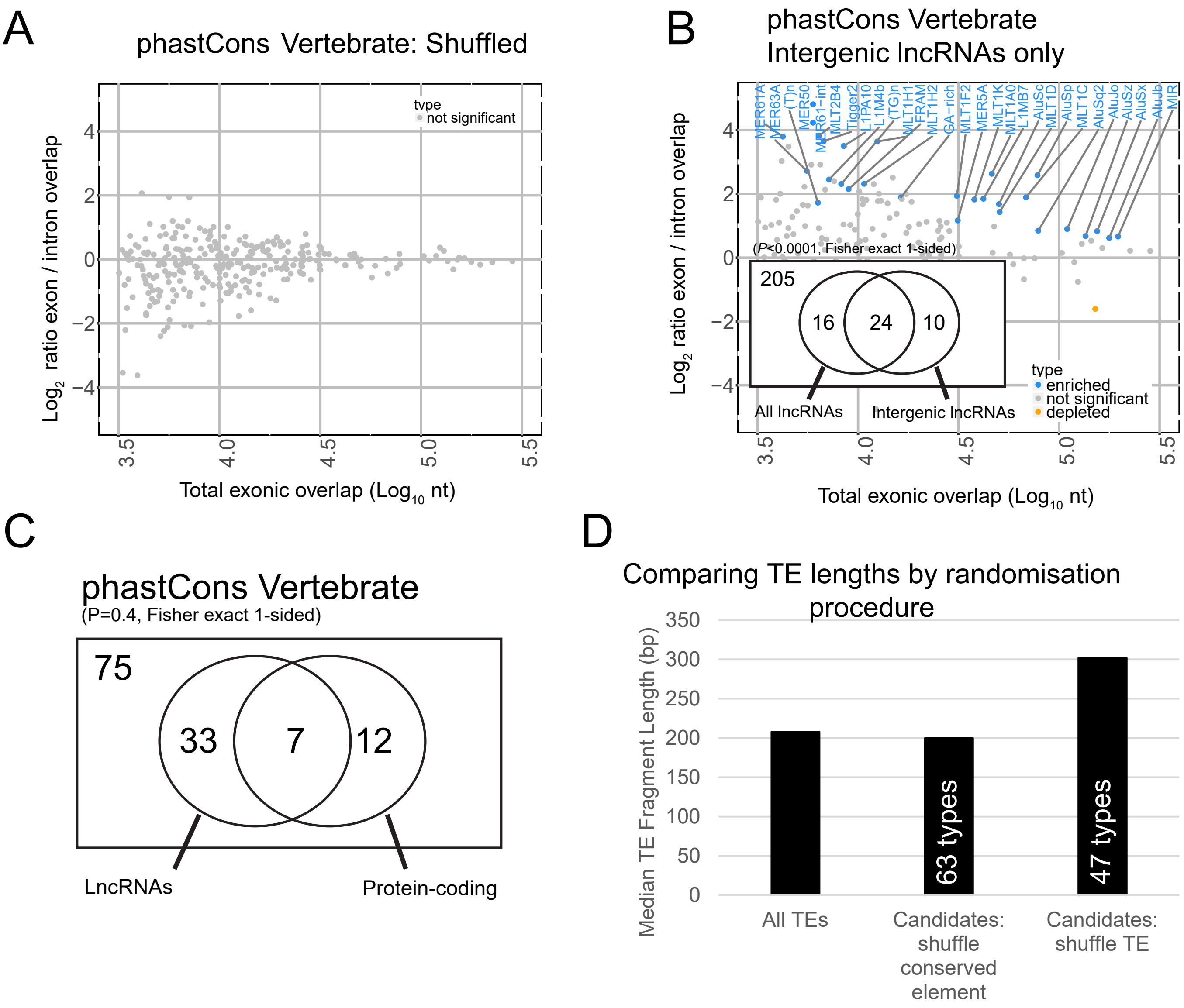


**Supplemental Figure S2:** (A) Figure shows, for every TE type, the ratio of exonic/intronic relative overlap by a shuffled set of phastCons Vertebrate elements. Significance was estimated by randomisation as in Figure 3, and here no significant TEs were identified. (B) Similar to (A), except that overlaps were performed using true phastCons Vertebrate conserved elements, and only the subset of intergenic lncRNA gene loci were included, ie those not overlapping a protein-coding locus. Blue indicates enriched TE types, as estimated by randomisation with an uncorrected p-value < 0.001. The inset shows the overlap of significant TE types between this analysis (right side) and the equivalent analysis using unfiltered lncRNAs (left, Supplemental Figure 1C). (C) Comparison of candidate RIDLs in lncRNAs and protein-coding genes, identified by overlap with phastCons Vertebrate elements. Numbers indicate TE types. (D) Comparison of lengths of significantly-conserved, exonically-overlapping TEs identified by two different randomisation procedures: shuffling conserved region locations, holding TEs constant (method used in Results); shuffling TE locations, holding conserved regions constant. For comparison, the lengths of the entire set of exonic TEs used in the study are shown ("All TEs"). Note that lengths are calculated using the entire TE fragment, and not just the portion overlapping the exon.
