## Supplementary material for "Ancient exapted transposable elements promote nuclear enrichment of human long noncoding RNAs"

Supplemental Figure S3

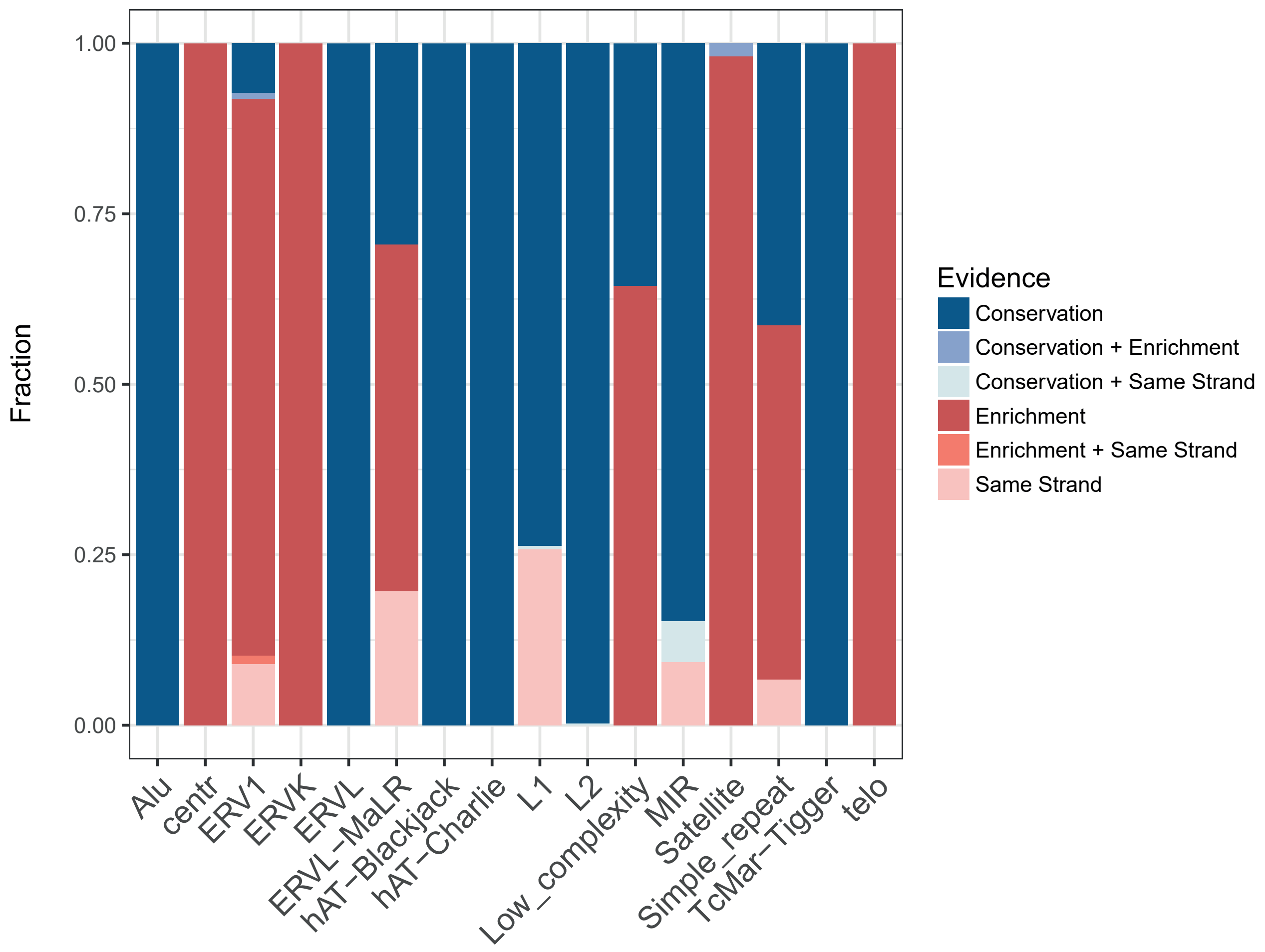


**Supplemental Figure S3:** Proportional frequencies of RIDLs identified by evidence type.
