## Supplementary material for "Ancient exapted transposable elements promote nuclear enrichment of human long noncoding RNAs"

Supplemental Figure S4

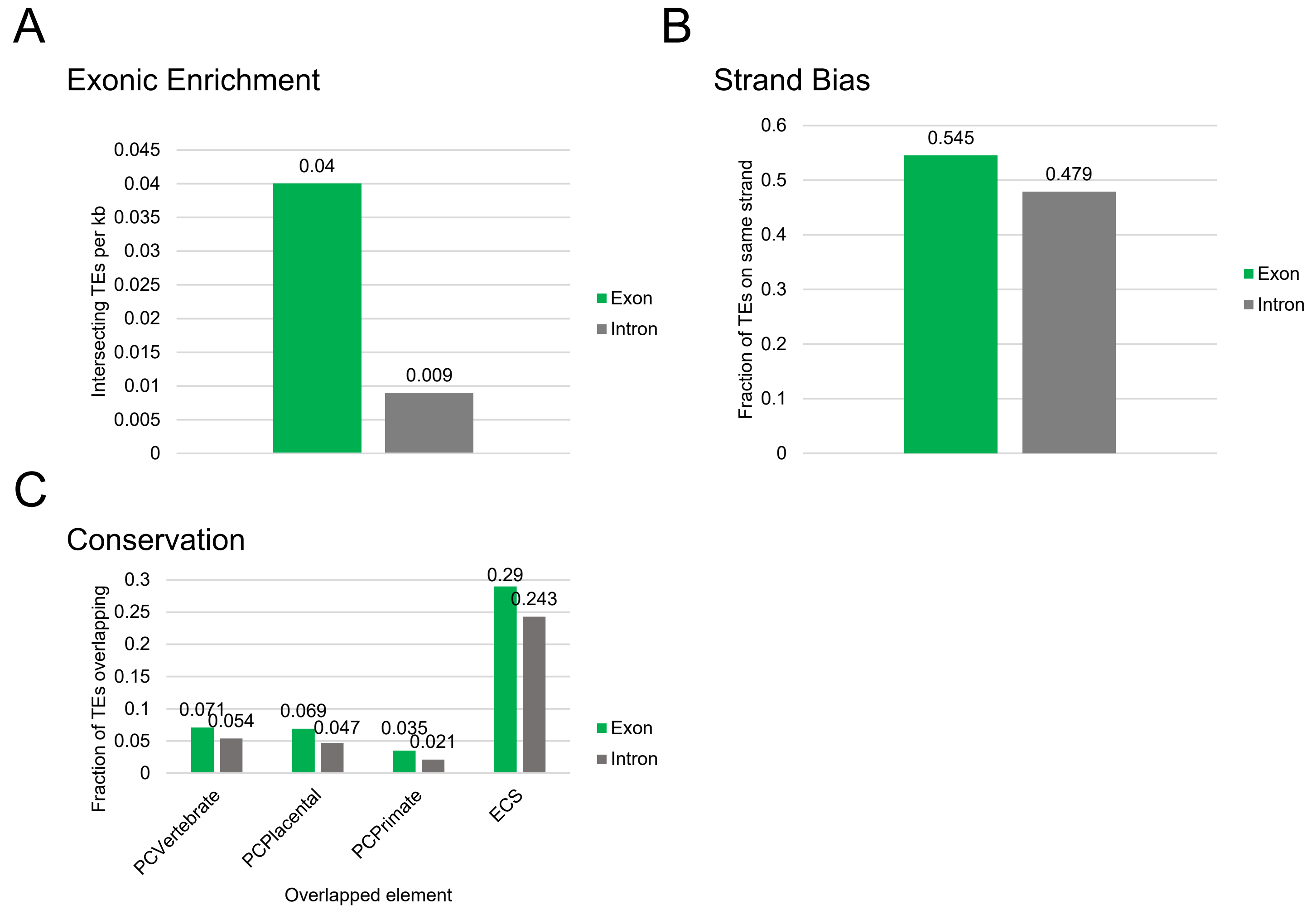


**Supplemental Figure S4:** Estimating true positive rates for RIDL predictions. Figures show fractions of TEs broken down by exonic and intronic cases. Only TEs from significant types for each evidence source, as outlined in Materials and Methods and shown in Figure D, are considered. We interpret the value for introns to represent an upper limit of the false discovery rate, and the difference between exon and intron to represent the number of true positive predictions. (A) The number of TEs from 20 types defined to be exonically enriched, which overlap either exons or introns of lncRNAs. (B) For RIDLs defined to have strand bias, the fraction of instances lying on the same strand as host lncRNA. (C) For RIDLs with evidence of evolutionary conservation, the fraction of instances that overlap the indicated conserved elements.
