## Supplementary material for "Ancient exapted transposable elements promote nuclear enrichment of human long noncoding RNAs"

Supplemental Figure S5

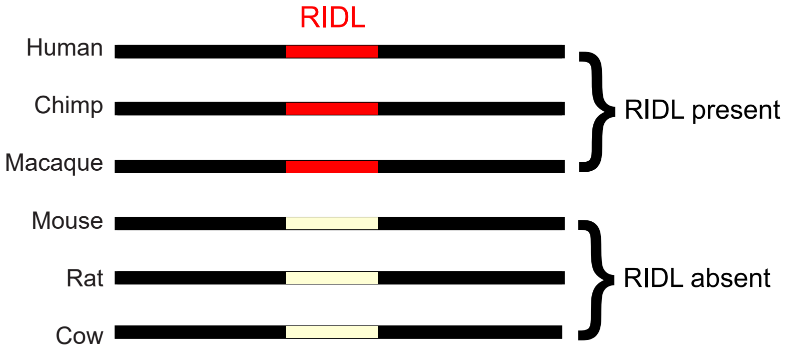


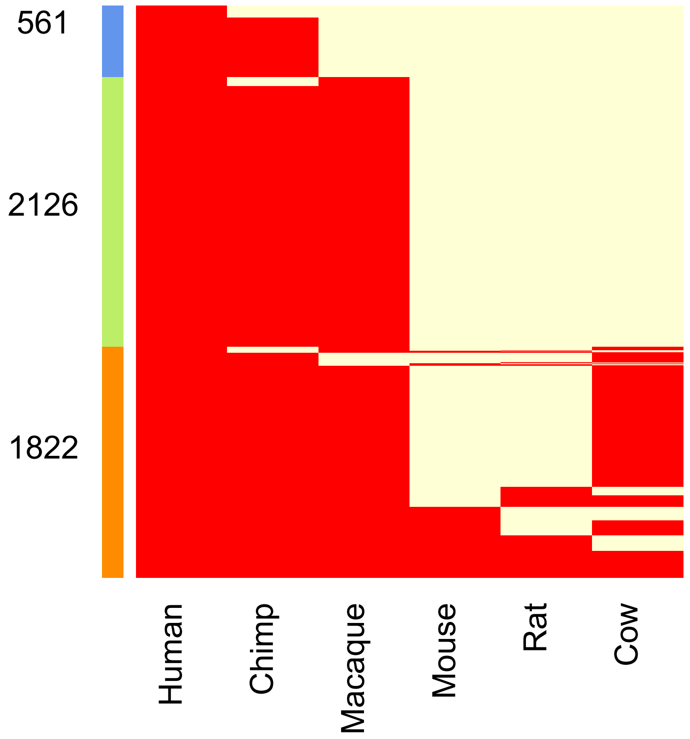


**Supplemental Figure S5:** Inferring the evolutionary age of RIDLs using 6-mammal alignments. Rows represent human RIDLs, columns represent species. Cells are coloured red when an orthologue is detected, otherwise in light yellow. Numbers represent the count of RIDLs conserved amongst Great Apes (blue), Primates (green) and Mammals (orange).
