## Supplementary material for "Ancient exapted transposable elements promote nuclear enrichment of human long noncoding RNAs"

### Supplemental Figure S6

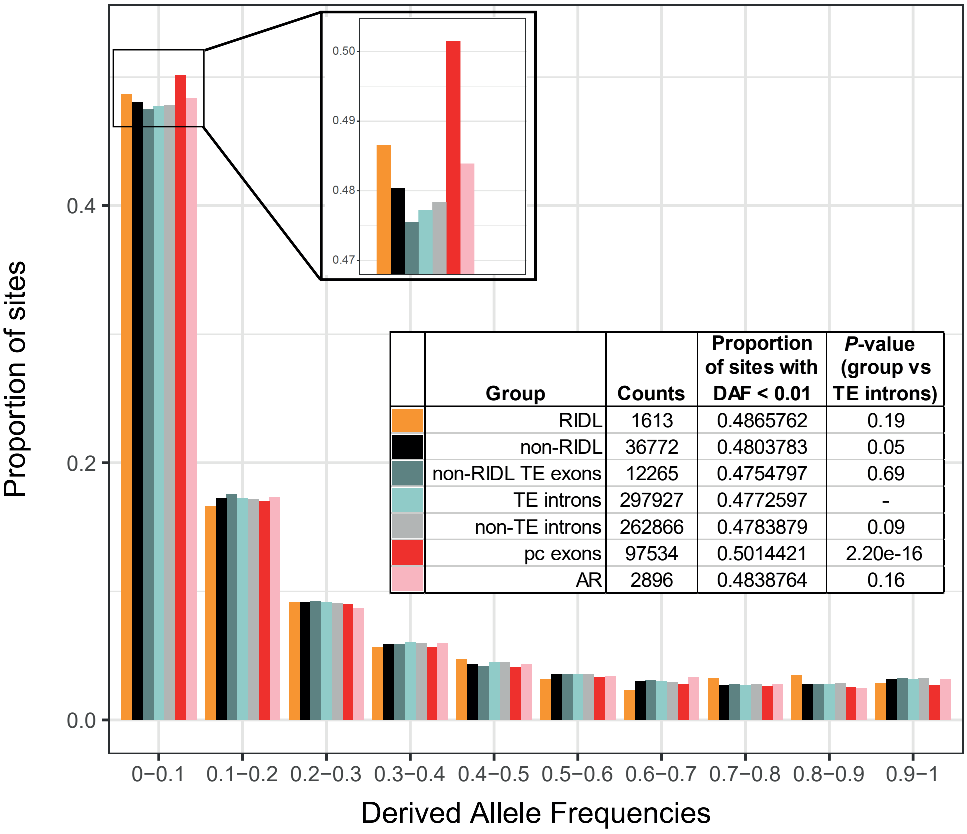


| | Group | Counts | Proportion of sites with DAF < 0.01 | *P*-value (group vs TE introns) |
|---|---|---|---|---|
| | RIDL | 1613 | 0.4865762 | 0.19 |
| | non-RIDL | 36772 | 0.4803783 | 0.05 |
| | non-RIDL TE exons | 12265 | 0.4754797 | 0.69 |
| | TE introns | 297927 | 0.4772597 | - |
| | non-TE introns | 262866 | 0.4783879 | 0.09 |
| | pc exons | 97534 | 0.5014421 | 2.20e-16 |
| | AR | 2896 | 0.4838764 | 0.16 |

**Supplemental Figure S6:** Proportion of common SNPs falling in the different bins of derived allele frequencies (DAF) for every group described in the table (left column). Protein coding exons (pc exons) and ancestral repeats (AR) are shown as positive and negative controls, respectively. The table shows for every group the total SNPs counts (second column) with low-frequency alleles (DAF < 0.1), the proportion this represents (third column) and the *p*-value obtained from comparing each group to TE introns (taken as a background) (Fisher's exact test, 1-sided).
