## Supplementary material for "Ancient exapted transposable elements promote nuclear enrichment of human long noncoding RNAs"

Supplemental Figure S7

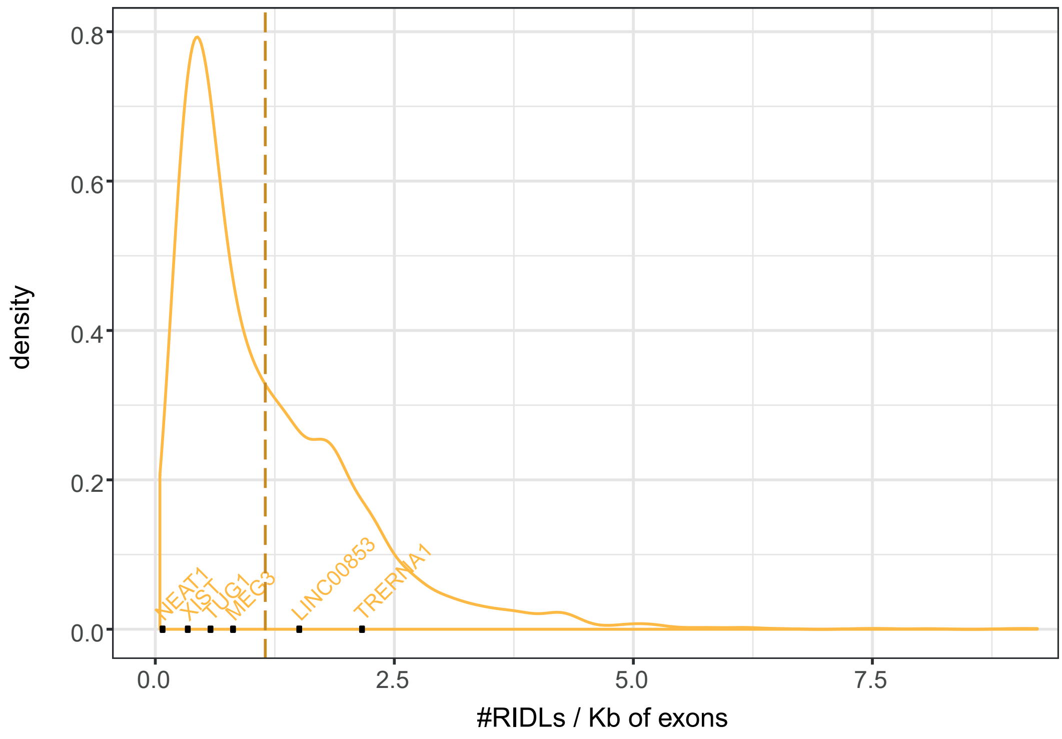


**Supplemental Figure S7:** Density distribution of RIDL-carrying lncRNAs based on the number of RIDLs per kb of exon. Dotted line indicates median. Selected known lncRNAs are indicated.
