## Supplementary material for "Ancient exapted transposable elements promote nuclear enrichment of human long noncoding RNAs"

Supplemental Figure S8

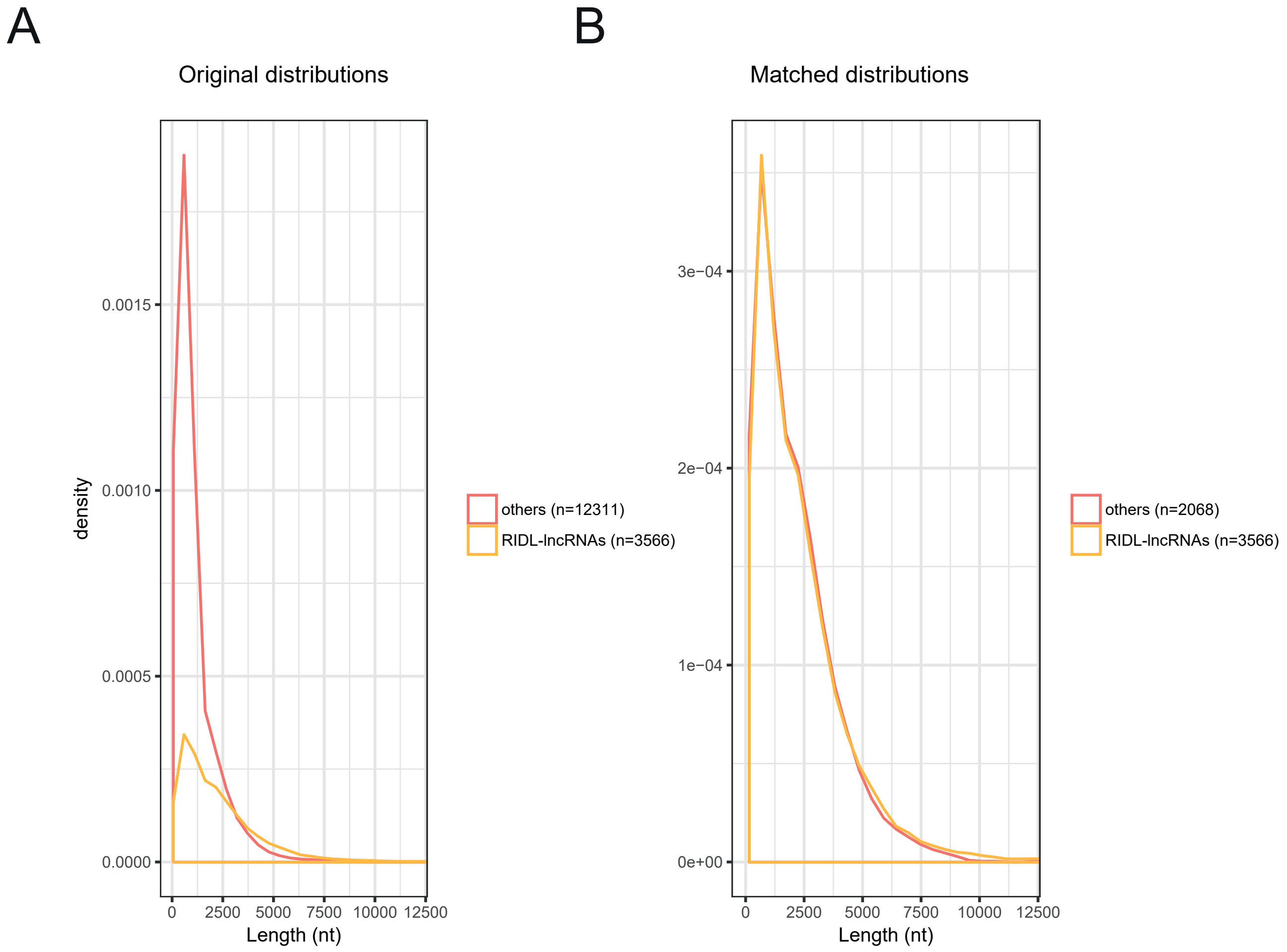


**Supplemental Figure S8:** Creating a length-matched sample of non-RIDL lncRNAs. (A) Exonic length distribution of RIDL-carrying (RIDL-lncRNAs) and non-RIDL ("others") GENCODE v21 lncRNAs, prior to sampling. (B) Exonic length distribution of RIDL-carrying lncRNAs and a length-matched sample of non-RIDL lncRNAs. The latter were used for all comparisons at gene level.
