## Supplementary material for "Ancient exapted transposable elements promote nuclear enrichment of human long noncoding RNAs"

Supplemental Figure S9

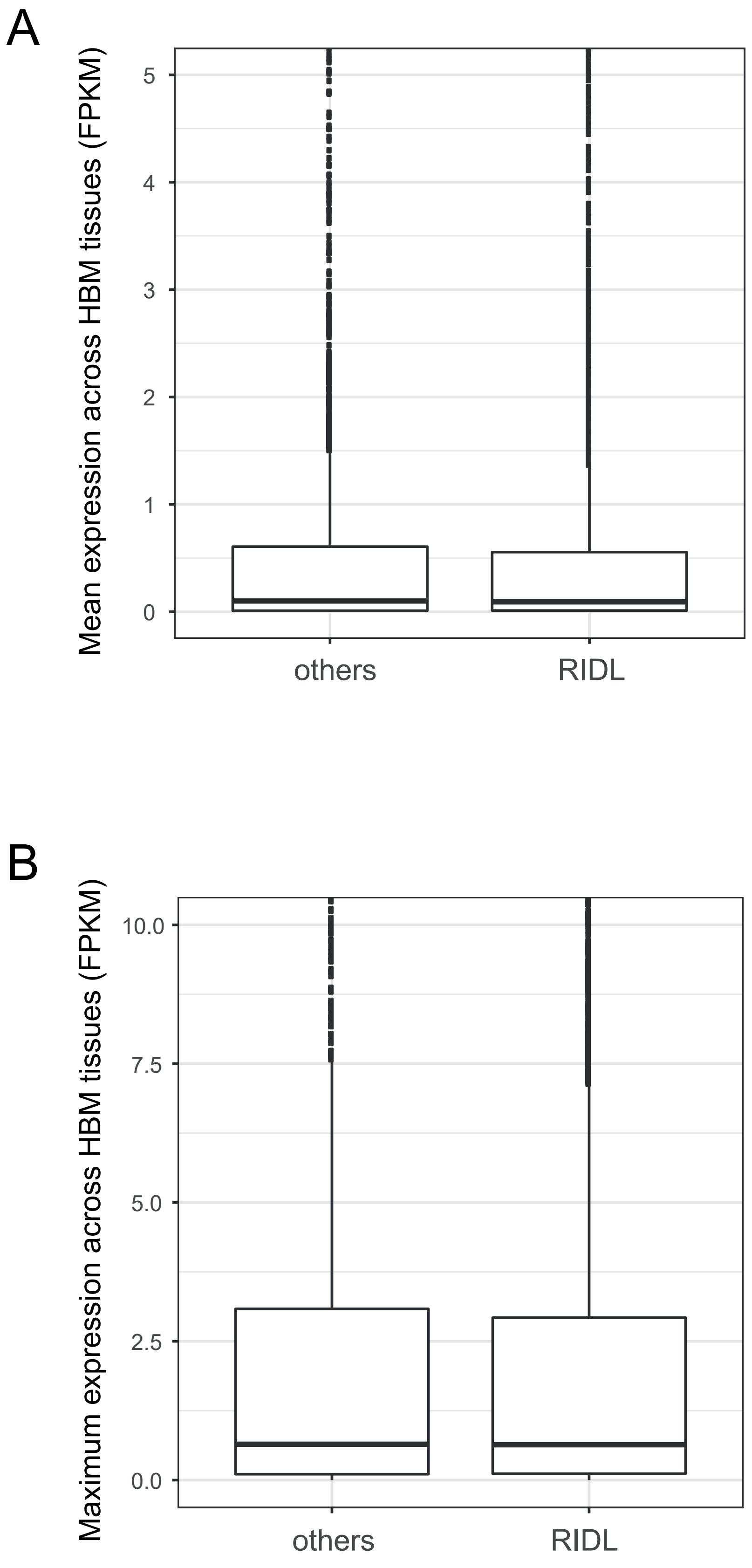


**Supplemental Figure S9:** (A) Distribution of mean expression across Human Body Map (HBM) tissues of RIDL-lncRNAs and other length-matched lncRNAs (others). (B) Same as (A) for maximum expression across HBM tissues.
