## Supplementary material for "Ancient exapted transposable elements promote nuclear enrichment of human long noncoding RNAs"

Supplemental Figure S10

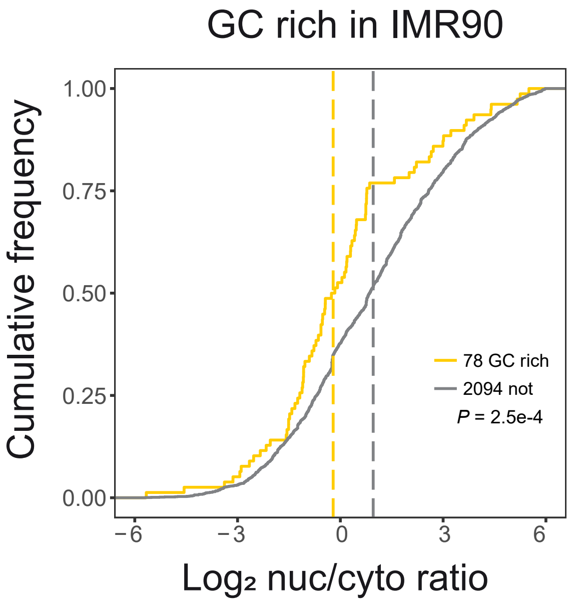


**Supplemental Figure S10:** Nuclear/cytoplasmic localization of GC-rich RIDL-containing lncRNAs in IMR90. Yellow represents lncRNAs carrying one or more copies of GC-rich RIDL elements, grey represents all other detected lncRNAs ("not"). Dashed lines indicate the median of each group. Significance was calculated using Wilcoxon test (P).
