## Supplementary material for "Ancient exapted transposable elements promote nuclear enrichment of human long noncoding RNAs"

Supplemental Figure S11

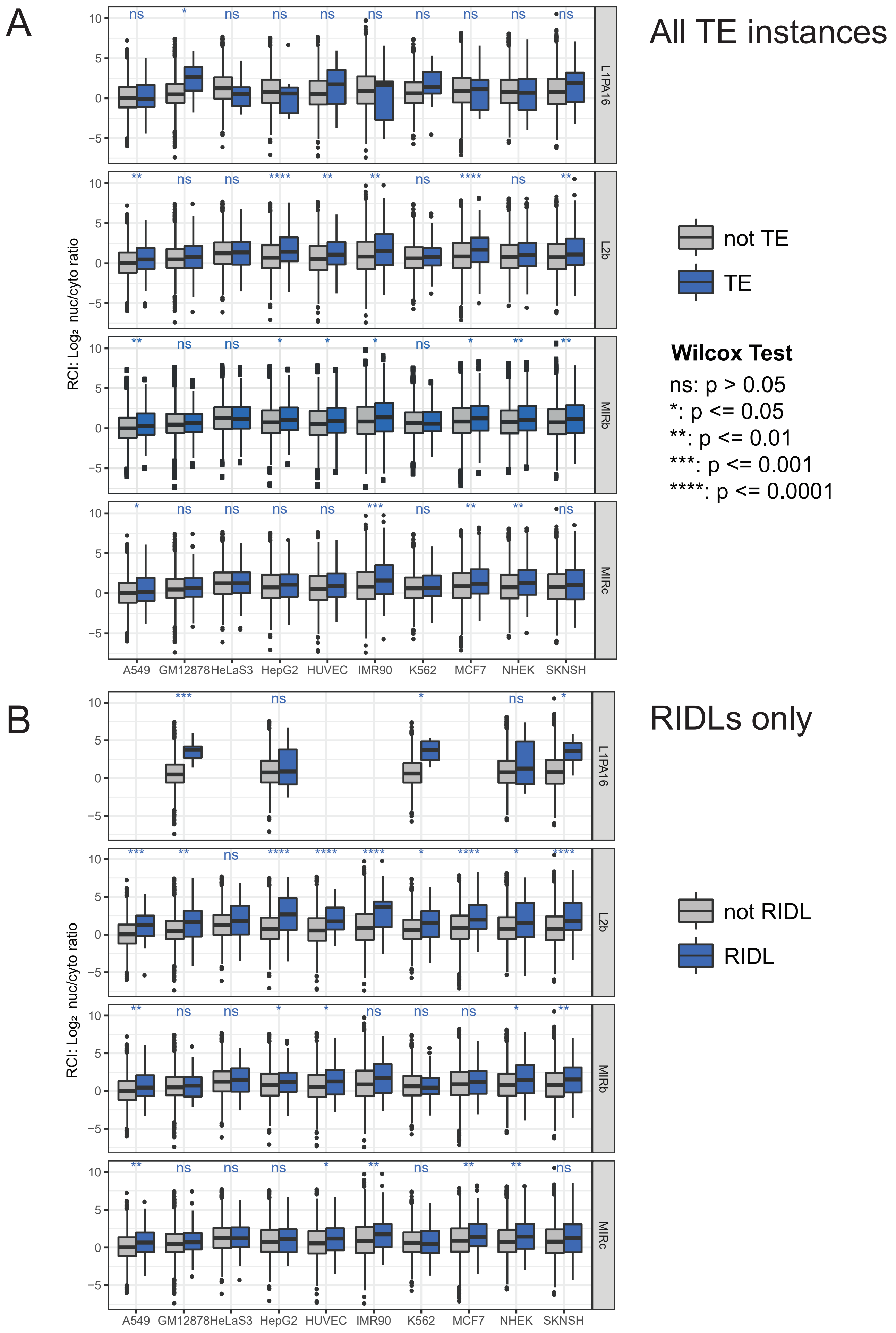


**Supplemental Figure S11:** Presence of L1PA16, L2b, MIRb and MIRc in lncRNA exons correlate with localisation, when considering both unfiltered TEs and RIDL sets. (A) For every cell line boxplots show RCI values of unfiltered TE-carrying lncRNAs versus non-TE-carrying lncRNAs (each panel contains information for a different TE type). (B) As for (A) but comparing RIDL-lncRNAs vs non-RIDL-lncRNAs.
