## Supplementary material for "Ancient exapted transposable elements promote nuclear enrichment of human long noncoding RNAs"

Supplemental Figure S12

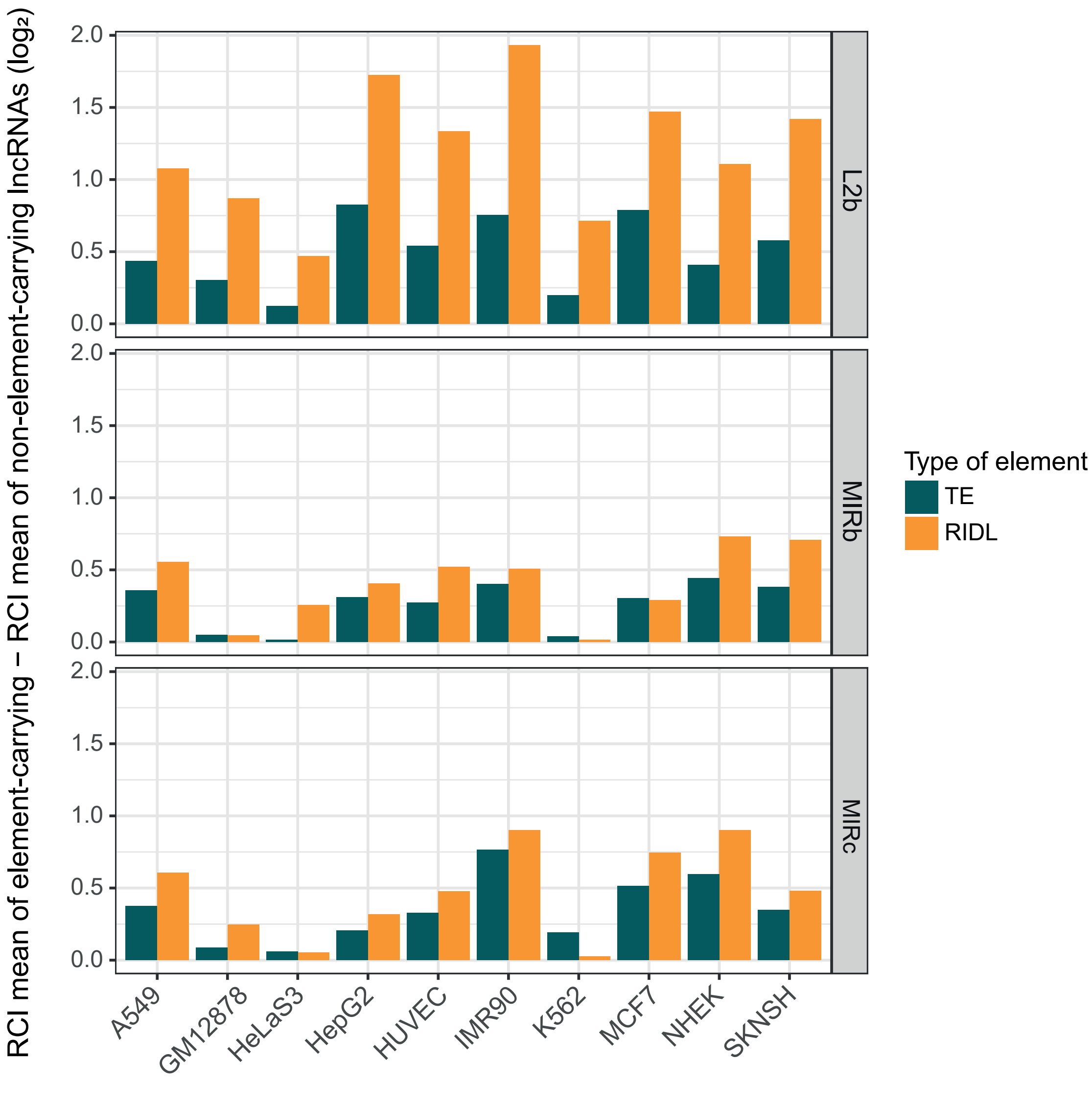


**Supplemental Figure S12:** RIDLs have a stronger effect on localisation than unfiltered TEs. Bars indicate the $log_2$ difference between the RCI (nuc/cyto ratio) of element-carrying lncRNAs and non-TE lncRNA.
