## Supplementary material for "Ancient exapted transposable elements promote nuclear enrichment of human long noncoding RNAs"

Supplemental Figure S13

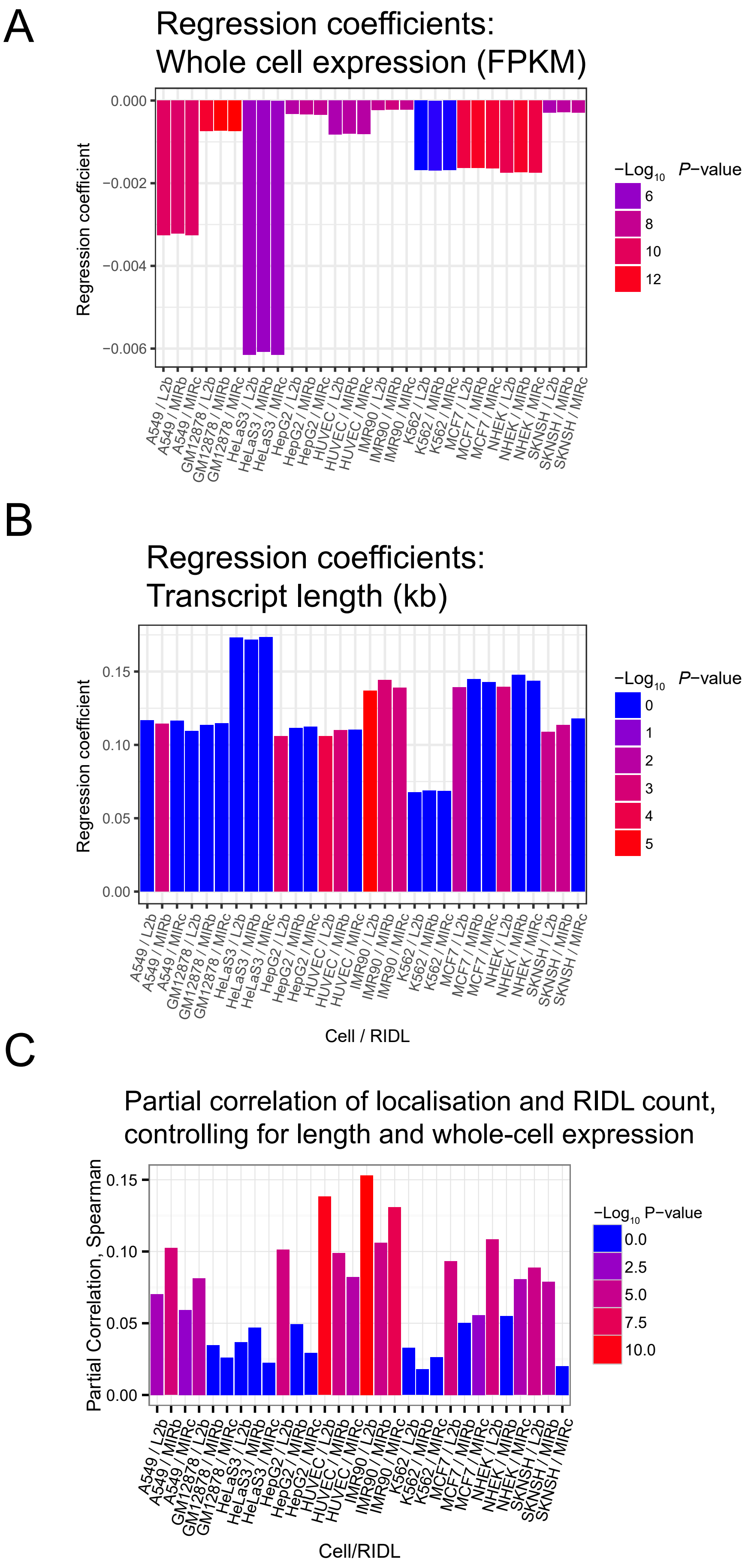


**Supplemental Figure S13:** (A) Regression coefficients for explanatory variable of whole-cell expression (in units of FPKM), in a linear model where dependent variable is lncRNA nuclear/cytoplasmic localisation (see Figure 5E and Methods). x-axis represents each RIDL/cell-line combination. Colours reflect estimated p-value for the association. (B) As for (A), but for explanatory variable of transcript length (in kb). (C) Partial correlation coefficients (Spearman) of $\log_2$ nuclear/cytoplasmic localisation and RIDL count, controlling for length and whole-cell expression. x-axis represents each RIDL/-cell-line combination. Colours reflect estimated p-value for the association.
