## Supplementary material for "Ancient exapted transposable elements promote nuclear enrichment of human long noncoding RNAs"

Supplemental Figure S14

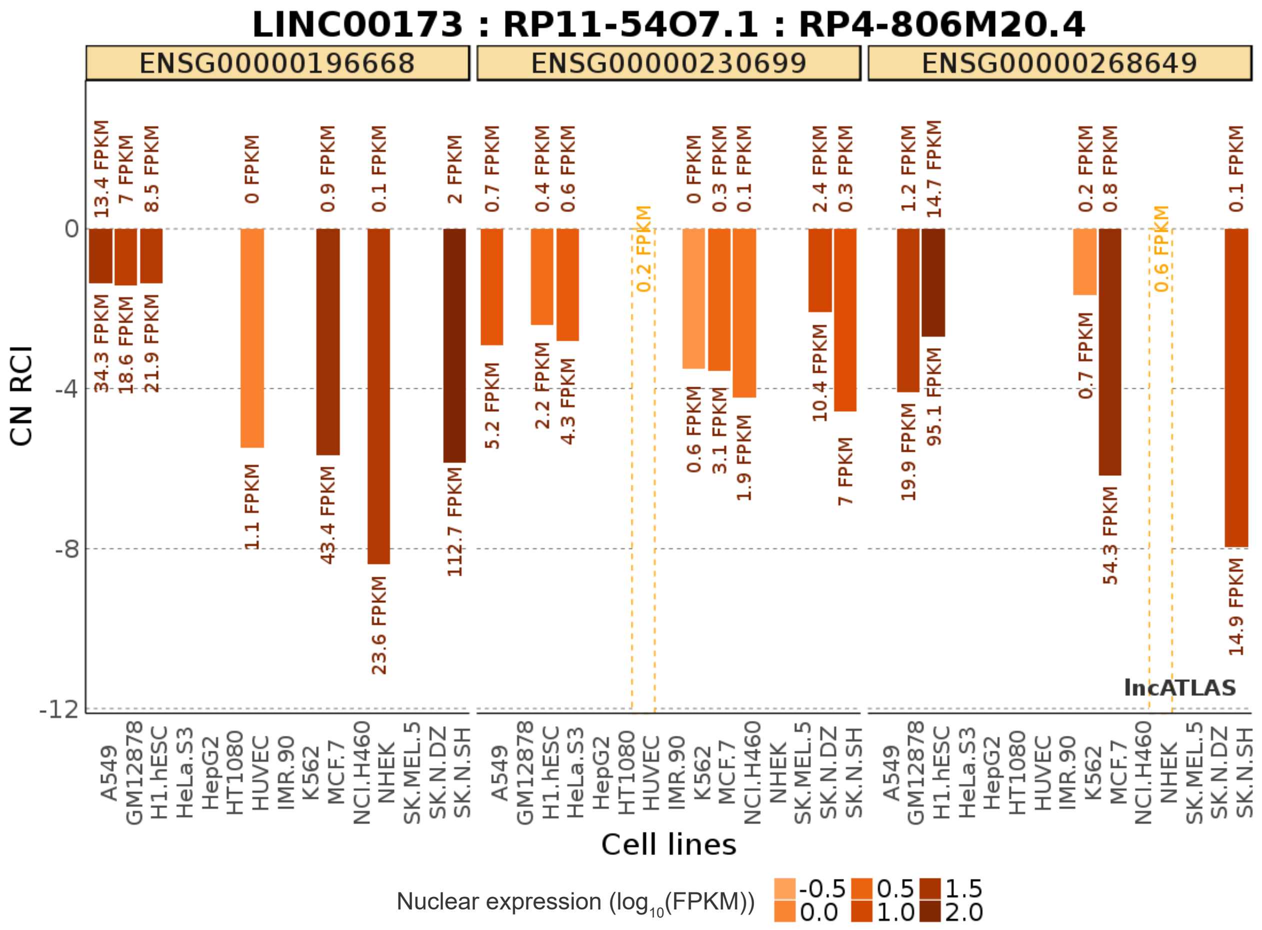


**Supplemental Figure S14:** Bars indicate log2-transformed cytoplasmic/nuclear expression ratios (CN RCI) in 15 cell lines for three candidate genes. Colours represent the total nuclear expression of each gene in each cell line. Barplot generated by LncATLAS webserver.
