## Supplementary material for "Ancient exapted transposable elements promote nuclear enrichment of human long noncoding RNAs"

Supplemental Figure S15

A

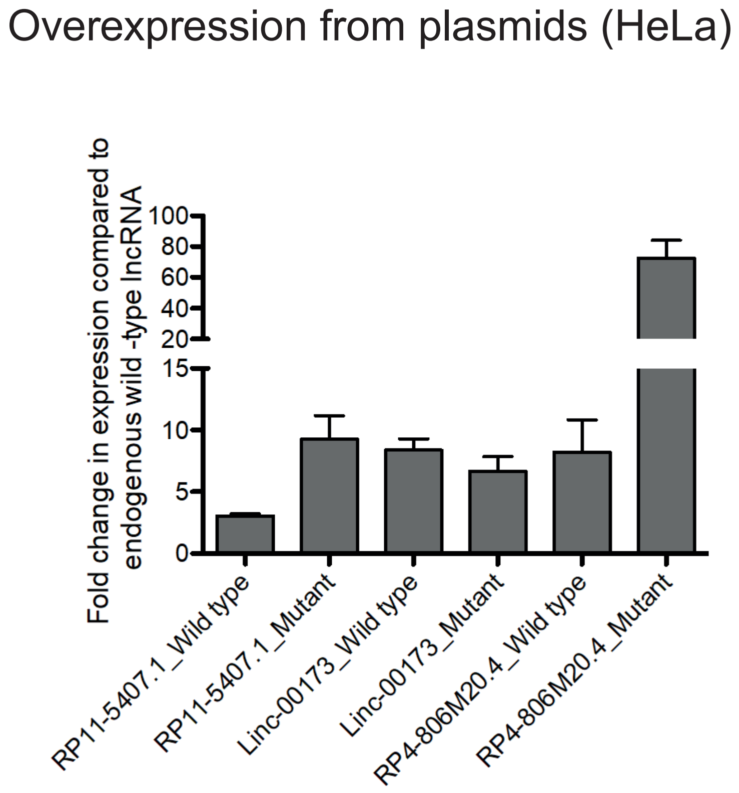


B

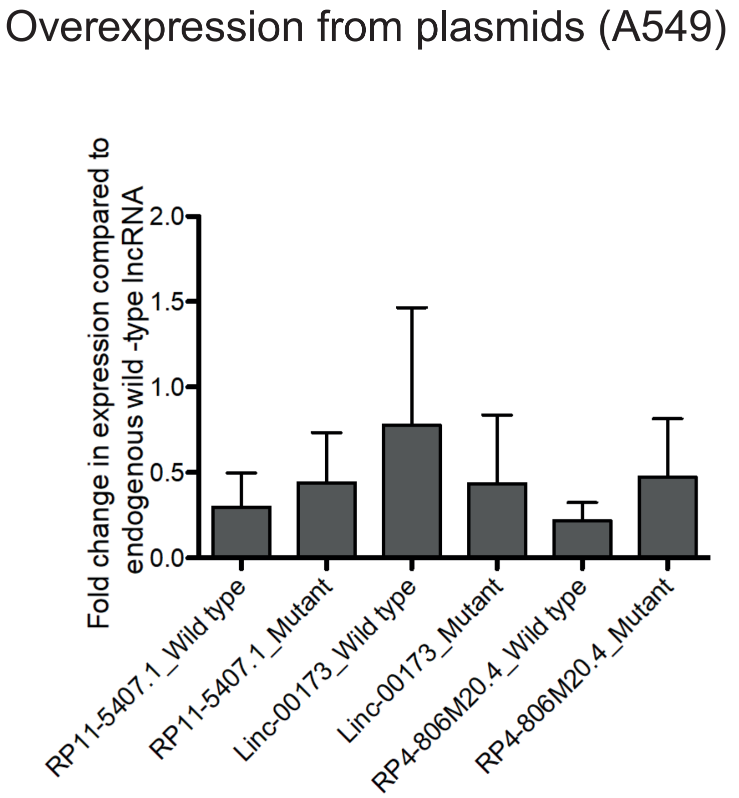


**Supplemental Figure S15:** Estimating overexpression of transfected lncRNA candidates (wild-type and mutant). Data are shown for the mean of four biological replicates. y axis represents the fold change of transfected gene level (normalised to GAPDH) compared to the endogeneous level (normalised to GAPDH) measured from untransfected cells. (A) HeLa cells, (B) A549 cells.
