## Supplementary material for "Ancient exapted transposable elements promote nuclear enrichment of human long noncoding RNAs"

Supplemental Figure S16

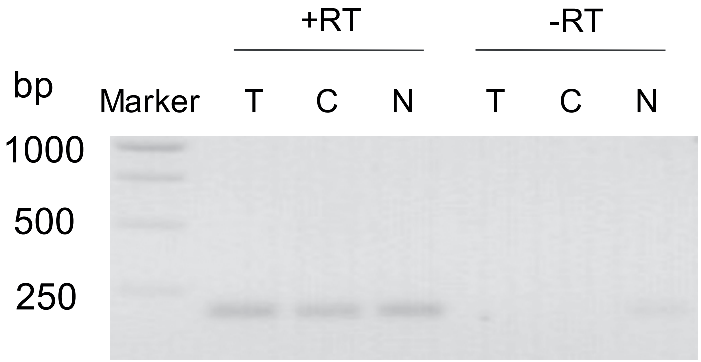


**Supplemental Figure S16:** DNA contamination check in RNA Isolated from different subcellular fractions. Total RNA was isolated from different subcellular fractions using RNA isolation kit as described in Materials and Methods. Equal amount of RNAs with (+RT) and without (-RT) reverse transcription, were subjected to PCR (40 cycles) with an exonic primer. The reaction products were electrophoresed on a 1.5 % agarose gel stained with SYBR safe. The T, C, N represent the RNA isolated from total, cytoplasmic and nuclear fractions respectively.
