## Supplementary material for "Ancient exapted transposable elements promote nuclear enrichment of human long noncoding RNAs"

Supplemental Figure S18

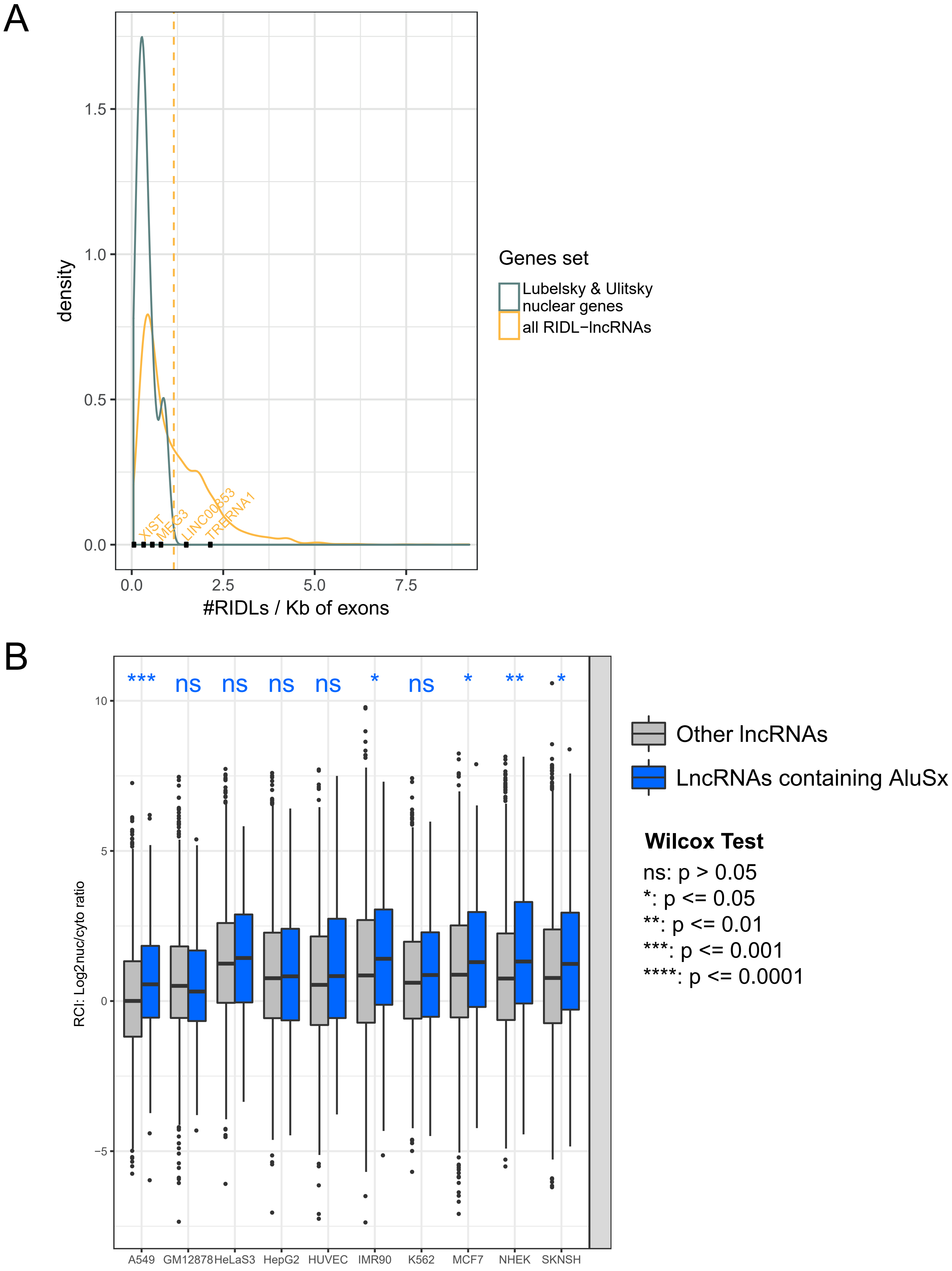


**Supplemental Figure S18:** Additional candidate lncRNA data for A549 cells. (A) Expression data for three candidate lncRNAs calculated using RNAseq data. (B) Validation of nuclear/cytoplasmic localisation of candidate genes in wild-type cells by qRT-PCR.
