## Supplementary material for "Ancient exapted transposable elements promote nuclear enrichment of human long noncoding RNAs"

Supplemental Figure S19

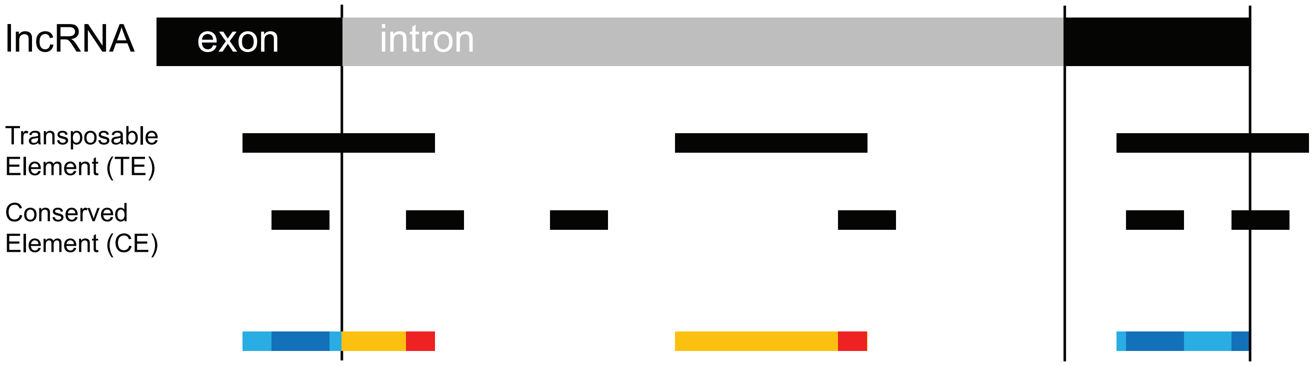


$$\text{Relative Exon Intron Conservation (REIC)} = \frac{\left(\dfrac{\text{Conserved exonic TE nts}}{\text{Conserved exonic TE nts} + \text{Non-conserved exonic TE nts}}\right)}{\left(\dfrac{\text{Conserved intronic TE nts}}{\text{Conserved intronic TE nts} + \text{Non-conserved intronic TE nts}}\right)}$$

**Supplemental Figure S19:** Overview of evolutionary analysis method
